# Direct Prediction of Intrinsically Disordered Protein Conformational Properties From Sequence: Version 3 [2023-05-28]

**DOI:** 10.1101/2023.05.08.539824

**Authors:** Jeffrey M. Lotthammer, Garrett M. Ginell, Daniel Griffith, Ryan J. Emenecker, Alex S. Holehouse

## Abstract

Intrinsically disordered regions (IDRs) are ubiquitous across all domains of life and play a range of functional roles. While folded domains are generally well-described by a single 3D structure, IDRs exist in a collection of interconverting states known as an ensemble. This structural heterogeneity means IDRs are largely absent from the PDB, contributing to a lack of computational approaches to predict ensemble conformational properties from sequence. Here we combine rational sequence design, large-scale molecular simulations, and deep learning to develop ALBATROSS, a deep learning model for predicting IDR ensemble dimensions from sequence. ALBATROSS enables the instantaneous prediction of ensemble average properties at proteome-wide scale. ALBATROSS is lightweight, easy-to-use, and accessible as both a locally installable software package and a point-and-click style interface in the cloud. We first demonstrate the applicability of our predictors by examining the generalizability of sequence-ensemble relationships in IDRs. Then, we leverage the high-throughput nature of ALBATROSS to characterize emergent biophysical behavior of IDRs within and between proteomes.

**Update from previous version:**

- This preprint reports an updated version of the ALBATROSS network weights trained on simulations of over 42,000 sequences.
- In addition, we provide new colab notebooks that enable proteome-wide IDR prediction and annotation in minutes.
- All conclusions and observations made in versions 1 and 2 of this manuscript remain true and robust.

## INTRODUCTION

Intrinsically disordered proteins and protein regions (IDRs) make up an estimated 30% of most eukaryotic proteomes and play a variety of roles in molecular and cellular function^1–4^. Although folded domains are often well-described by a single (or small number of) three-dimensional (3D) structures, IDRs are defined by extensive conformational heterogeneity. This means they exist in a conformational ensemble - a collection of rapidly interconverting states that prohibits structural classification by any single reference structure^5, 6^. This heterogeneity challenges many experimental, computational, and conceptual approaches developed for folded domains, necessitating the application of polymer physics to describe, classify, and interpret IDRs in a variety of contexts^7–18^.

Although IDRs are defined by the absence of a defined folded state, they are not “unstructured”^19^. The same chemical moieties that drive protein folding and enable molecular recognition in folded domains are also found within IDRs. As such, while folded domains subscribe to a sequence-structure relationship, IDRs have an analogous sequence-ensemble relationship^19^. Over the last fifteen years, there has been a substantial effort to decode the mapping between IDR sequence and conformational properties, the so-called ‘sequence-ensemble relationship ^5, 12, 19–35^.

IDR conformational properties can be local or global. Local conformational properties typically involve transient secondary structure, especially transient helicity^36^. Global conformational properties report on ensemble-average dimensions - that is, the overall size and shape that the ensemble occupies^5, 19, 36, 37^. Two common properties measured by both experiment and simulation are the radius of gyration (R_g_) and end-to-end distance (R_e_). The R_g_ reports on the volume an ensemble occupies, while the R_e_ reports on the average distance between the first and the last residue. Ensemble shape can be quantified in terms of asphericity, a parameter that lies between 0 (sphere) and 1 (prolate ellipsoid), and reports on how spherical an ensemble is. While R_e_, R_g_, and asphericity are relatively coarse-grain, they can offer insight into the molecular conformations accessible to an IDR, as well as provide hints at the types of intramolecular interactions which may also be relevant for intermolecular interactions (especially in the context of low-complexity sequences)^23, 38, 39^.

An *in vitro* assessment of sequence-ensemble relationships involves expression, purification, and measurement of ensemble properties using various biophysical techniques^16, 40, 41^. The experimental methods commonly used to study conformational properties include single-molecule fluorescence spectroscopy, nuclear magnetic resonance (NMR) spectroscopy, and small angle X-ray scattering (SAXS) ^16, 40–42^. While powerful, all three of these approaches can be technically demanding, necessitate access to specific instrumentation, and in the case of NMR and SAXS, require relatively high concentrations of protein. Beyond *in vitro* assessment, integrating all-atom simulations with biophysical measurements has proven invaluable in obtaining a holistic description of sequence-ensemble relationships, yet these integrative studies can also be challenging ^22, 23, 25, 28, 43–47^. As such, obtaining insight into sequence-specific conformational biases for disordered proteins is often challenging for groups with a limited background in molecular biophysics.

Recent efforts have led to a marked improvement in the accuracy of coarse-grained force fields for disordered protein simulations ^48–53^. In particular, simulations performed with the CALVADOS and Mpipi force fields offer robust predictions of global conformational properties for disordered proteins. However, setting up, running, and analyzing molecular simulations necessitate a level of expertise and resources beyond many (arguably most) research groups. As such, the democratization of exploring sequence-to-ensemble relationships in disordered proteins demands easy-to-use tools that are readily accessible (i.e., available in a web browser without any hardware constraints).

Here, we address this gap by developing a rapid and accurate predictor for disordered protein global dimensions from sequences. We do this through a combination of rational sequence design, large-scale coarse-grained simulations, and deep learning (**Fig. 1A**). The resulting predictor (ALBATROSS; <u>A</u> deep-<u>L</u>earning <u>B</u>ased <u>A</u>pproach for predic<u>T</u>ing p<u>R</u>operties <u>O</u>f di<u>S</u>ordered protein<u>S</u>) not only pushes the boundaries of acronym development but provides a means to predict IDR global dimensions (R_g_, R_e_, asphericity, and apparent polymer scaling exponent) directly from sequence.

**Figure 1.**
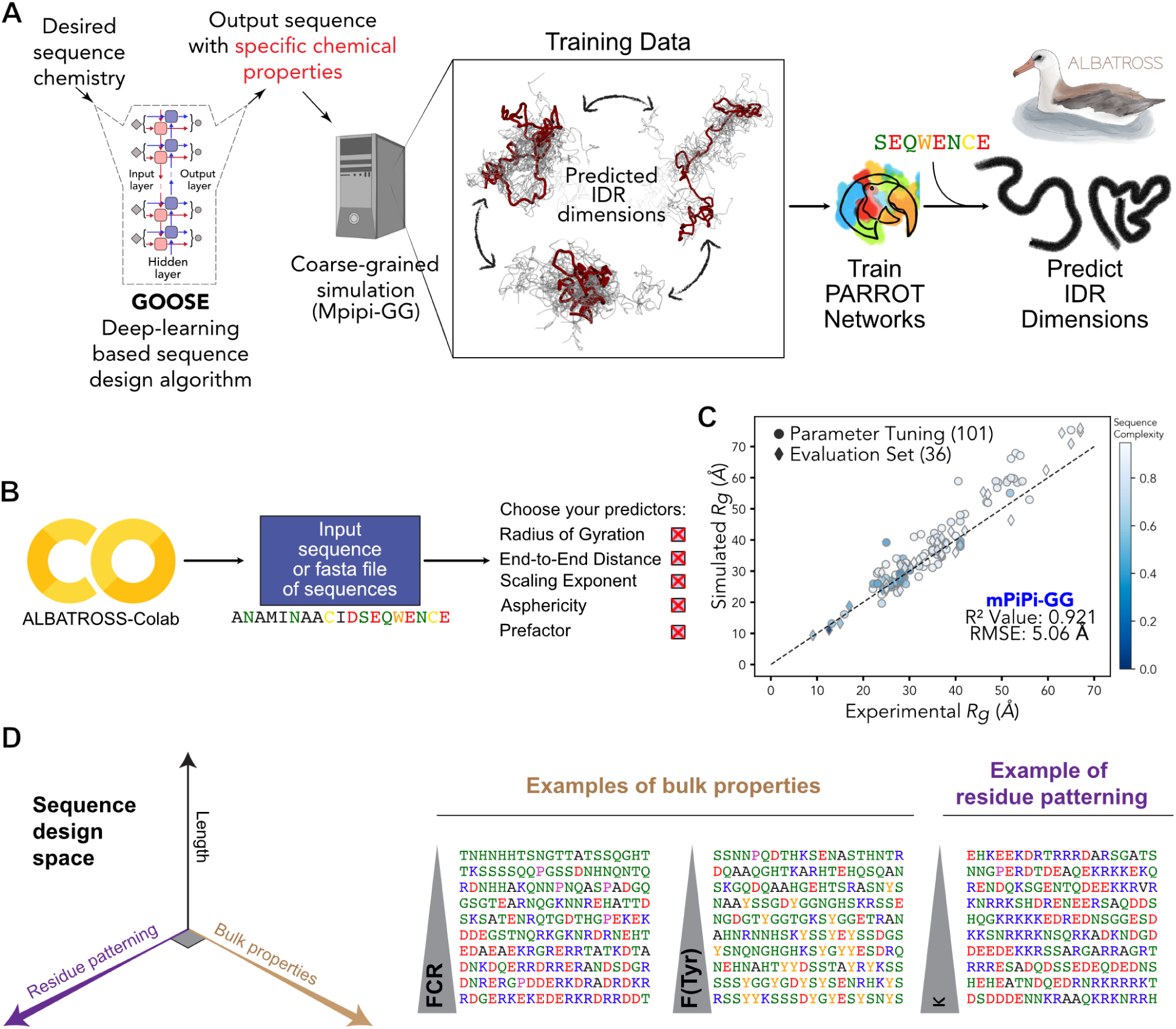
ALBATROSS is a deep-learning framework for predicting sequence-dependent IDR ensemble properties. A) Sequence design and simulation approach to generate training data for ALBATROSS networks. The Python package GOOSE is used to generate synthetic IDRs across a diverse area of sequence space. Coarse-grained molecular dynamics simulations are performed for each sequence to generate labeled data for downstream deep neural network training and validation. **B)** ALBATROSS is implemented as a point-and-click style interface on Google Colaboratory with support for CPU and GPU inference. The user simply specifies the amino acid sequence or a fasta file of amino acid sequences and then selects the predictions they would like to perform. **C)** Correlation and RMSE between the Mpipi-GG force field and a curated set of 137 experimental radii of gyration. 101 sequences (circles) were used for validating the Mpipi-GG force field, and 36 were new sequences held-out during parameter fitting (diamonds). The blue color gradient signifies the Wootton-Federhen complexity of the sequence. D) Rational sequence design scheme. GOOSE was used to design sequences that titrated along different protein sequence parameter axes: length, residue patterning, and bulk amino acid properties. The fraction of charged residues and the fraction of tyrosine residues are examples of bulk properties. An example of residue patterning is the sequence parameter Κ which describes the asymmetry of positively and negatively charged amino acids in a sequence,

ALBATROSS was developed with ease of use and portability in mind; no specific hardware is required, and predictions can be performed on either CPUs or GPUs. We provide both a locally-installable implementation of ALBATROSS as well as point-and-click Google Colab notebooks that enable predictions to be performed on 30-60 sequences per second on a CPU and thousands of sequences per second on a GPU (**Fig. 1B**). In this work, we use ALBATROSS to demonstrate the generality of core sequence-ensemble relationships identified by foundational prior work, as well as assess general conformational biases observed at proteome-wide scales. Lastly, we propose that local conformational behavior offers a route to discretize IDRs into conformationally-distinct subdomains.

By combining ALBATROSS with recent improvements to our state-of-the art disorder predictor (metapredict V2-FF), we also provide the ability to predict and annotate the entire set of IDRs for a given proteome (the IDR-ome) in seconds-to-minutes. This advance opens the door for large-scale unstructural bioinformatics and, importantly, makes these predictions easy for anyone with an internet connection.

As a final note, while we rigorously validate the accuracy of ALBATROSS against both simulated and experimental data, we do not see it as a replacement for well-designed simulation or experimental studies. Instead, our goal is for ALBATROSS to estimate emergent biophysical properties from IDR-encoded sequence chemistry to aid in hypothesis generation as well as the interpretation and design of experiments.

## METHODS

The overall approach for developing ALBATROSS involved several steps. First, we generated a library of synthetic disordered proteins that systematically titrated across compositional space using our artificial disordered protein design package GOOSE^54^. Next, we fine-tuned the Mpipi force field, making small changes to the previously published parameters to address minor shortcomings, leading to a version we refer to as Mpipi-GG^49^. We then performed simulations of the synthetic training sequences using Mpipi-GG and calculated ensemble-average parameters^55^. Finally, we trained bidirectional recurrent neural networks with long short-term memory cells (LSTM-BRNNs) to map between amino acid sequence and simulation-derived ensemble-average parameters^56^. Network weights, along with software to perform sequence-ensemble predictions, were then packaged into our sequence analysis package SPARROW and via an easy-to-use Google Colab notebook^57^.

### ALBATROSS Training Sequence Library Design

Using the IDR design package GOOSE, we assembled a library of chemically diverse synthetic disordered proteins (https://github.com/idptools/goose)^54^. Sequences varied charge, hydropathy, and charge patterning, as well as titrated across the amino acid composition. All sequences generated were between 10 and 750 residues in length. In total, we generated 23,127 disordered protein sequences across a diverse sequence space (see *Supplementary Information*). In addition to the synthetic sequence library, we curated a set of 19,075 naturally occurring IDRs by randomly sampling disordered proteins ranging in length from 10-750 residues from one of each of the following proteomes: *Homo sapien, Mus musculus, Dictyostelium discoideum, Escherichia coli, Drosophilia melanogaster, Saccharomyces cerevisiae, Neurospora crassa, Schizosaccharomyces pombe, Xenopus laevis, Caenorhabditis elegans, Arabidopsis thaliana, and Danio rerio.* All annotated IDRs from the aforementioned proteomes are available at https://github.com/holehouse-lab/shephard-data/tree/main/data/proteomes.

### ALBATROSS Validation Sequence Library Design

To prepare a test set to accurately assess the true generalization error for each of our ALBATROSS predictors, we randomly selected an additional set of 2,501 biological IDRs from one of the aforementioned proteomes. To ensure that any newly selected biological sequences were distinct from biological sequences seen during training, we leveraged CD-HIT with default parameters to remove sequences with >20% similarity^58^. In total, we were left with 2,306 biological IDRs in our test set. In addition, we also designed and simulated 3,731 synthetic disordered protein sequences. As before, these synthetic sequences varied charge, hydropathy, and charge patterning, as well as titrated across the amino acid composition. All sequences generated were between 10 and 750 residues in length. In total, the ALBATROSS test set we used to assess model accuracy consisted of 6,037 disordered protein sequences.

### Coarse-Grained Simulations

All reported simulations were performed with the LAMMPS simulation engine and either the newly parameterized Mpipi-GG or Mpipi (for comparison to Mpipi-GG) force fields^49, 59^. Initial disordered protein starting configurations were built by assembling beads as a random coil in the excluded volume limit. Each simulation was minimized for a maximum of 1000 iterations or until the force tolerance was below 1 x 10^-8^ (kcal/mol)/Å. All simulations were performed with 150 mM implicit salt concentration in the canonical (NVT) ensemble at a target temperature of 300 K. The simulation temperature was maintained with a weakly-coupled Langevin thermostat that is adjusted every 100 picoseconds, and an integration timestep of 20 femtoseconds for all production runs. Simulations were performed with periodic boundary conditions in a 500 Å^3^ cubic box. Output coordinates for each trajectory were saved every 2 nanoseconds. All simulations were initially equilibrated for 10 ns, and structures from this equilibration period were discarded. Production simulations of disordered sequences with less than 250 residues were performed for 6 µs, whereas sequences greater than 250 residues were simulated for 10 µs. In terms of LAMMPS simulation parameters, these settings reflect saving IDR conformations every 1 x 10^5^ simulation steps, discarding the first 5 x 10^5^ simulation steps as equilibration, and performing simulations for 3 x 10^8^ steps for short sequences and 1 x 10^9^ steps for long sequences. Simulation analysis was performed using SOURSOP and MDTraj ^55, 60^. We made extensive use of GNU parallel for simulation analysis^61^.

### Deep Learning

We leveraged Bidirectional Recurrent Neural Networks with Long Short-Term Memory cells (BRNN-LSTM) for all sequence-to-ensemble property prediction tasks with the flexible recurrent neural network framework PARROT^56^. We generated training, validation, and test data from coarse-grained simulations performed with the Mpipi-GG force field. Specifically, we developed predictors for the radius of gyration (R_g_), end-to-end distance (R_e_), and asphericity, along with the polymer scaling law prefactors and scaling exponents ^62, 63^.

Following previous PARROT network protocols, we employed a one-hot encoding scheme to translate the protein sequence data into numerical vectors amenable for deep neural network training. We used a training objective that sought to minimize an L1 loss function between the predictions and labeled data for each of the sequence-to-ensemble property predictors. For each of these prediction tasks, we performed a hyperparameter grid search with 5-fold cross validation for 500 epochs with PARROT (64% training, 16% validation, and 20% test). The set of hyperparameters that performed best on average across each fold were selected to train a final model with 80% of the data used as training data and 20% as validation data. Final network weights were chosen by selecting the epoch with the lowest validation loss across 750 epochs.

For each network, we chose a default learning rate of 0.001, and we performed a hyperparameters grid search over the following parameters: number of hidden layers (1 to 2), and a hidden dimension size (10 to 55). To evaluate the generalization error of our models on sequences relevant to biological function, we evaluated the most accurate networks for each predictor using the curated test set of 6,051 IDR sequences which consisted of both synthetic and naturally occurring IDRs.

### Disorder prediction

Disorder prediction (in this manuscript and in the associated notebooks) is provided through metapredict V2-FF ^64, 65^. Metapredict V2-FF is our newly implemented version of metapredict V2, which offers a 5-50x improvement in performance compared to metapredict V2 with no loss in prediction accuracy. V2-FF was developed specifically in the context of this manuscript for working with ALBATROSS, and enables proteome-wide prediction to be obtained in a reasonable (<10 min) timeframe.

### Bioinformatics

Proteome-wide bioinformatics was performed using SPARROW (https://github.com/idptools/sparrow) and SHEPHARD^66^. SPARROW is an in-development Python package for calculating IDR sequence properties, while SHEPHARD is a hierarchical analysis framework for annotating and analyzing large sets of protein sequences. IDRs and proteome data are available at https://github.com/holehouse-lab/shephard-data. Proteomes were obtained from UniProt^65, 67^. Normalized chain dimensions (normalized R_e_ and normalized R_g_) were calculated as the ALBATROSS-predicted R_e_ or R_g_ divided by the Analytical Flory Random Coil (AFRC)-derived R_e_ or R_g_. The AFRC is a model that reports on the sequence-specific chain dimensions expected if an IDR behaved as a Gaussian chain (i.e., a Flory scaling exponent of 0.5)^68^.

### Yeast homologous IDR analysis

Homologous yeast proteomes were obtained from the Yeast Genome Order Browser (YGOB)^69, 70^. Syntenic genes were used to identify homologous proteins, and these sequences were then aligned using Clustal Omega^71^. For all homolog sets where there were more than 10 total proteins and a *S. cerevisiae* protein present, this protein was segmented into folded and disordered domains using metapredict (V2)^64, 65^. This domain prediction was projected from the *S. cerevisiae* protein onto the multiple sequence alignment of all of the homologs, assigning any gapped regions between domains as IDRs. All sets of homologous IDRs where the *S. cerevisiae* IDR was 40 amino acids or longer were analyzed further. All IDRs belonging to one of these sets had predicted dimensions using ALBATROSS. Additionally, on each of these sets, sequence similarity of the aligned IDRs was calculated using the pyMSA (v0.5.1) package (https://github.com/benhid/pyMSA), using the BLOSUM62 scoring matrix. Both the SumOfPairs and StarScore metrics for evaluating MSA similarity were normalized by the number of aligned sequences and length of the alignment. A similar procedure was applied to the E1A linker sequences from^72^. Sequence features for the Yak1 and Spt2 IDR homologs were computed using SPARROW.

### ALBATROSS implementation and distribution

ALBATROSS is implemented within the SPARROW sequence analysis package (https://github.com/idptools/sparrow). In addition, a point-and-click style interface to ALBATROSS is provided via a stand-alone Google Colab notebook for both single-sequence and large-scale predictions of hundreds of sequences. If a FASTA file is uploaded and GPUs are selected, this notebook enables predictions for thousands of IDRs per second, facilitating in-browser proteome-wide analysis.

The notebook is available at: https://colab.research.google.com/github/holehouse-lab/ALBATROSS-colab/blob/main/example_notebooks/polymer_property_predictors.ipynb

For IDRs predicted from protein sequences at https://metapredict.net/, the predicted R_g_ and R_e_ are also returned instantaneously.

Finally, we provide a standalone notebook for predicting and annotating all IDRs in a proteome (i.e., IDR-ome construction): https://colab.research.google.com/github/holehouse-lab/ALBATROSS-colab/blob/main/idrome_c onstructor/idrome_constructor.ipynb

Specifically, IDR-ome construction combines predicting IDRs across an entire proteome with calculating IDR sequence properties and predicted IDR ensemble properties. With GPU-support on Google colab, this notebook enables the construction of the annotated human IDR-ome in ∼5-8 minute. Without GPU support, the same output is achieved in ∼7-10 minutes.

### Data and code availability

All of the data associated with the proteome-wide analysis presented in **Fig. 4** and **Fig. 5** are shared as SHEPHARD-compliant datafiles, and we encourage other groups to explore these predictions in the context of other protein annotations using SHEPHARD and the set of precomputed annotations provided therein^66^. All other data and code used for sequence analysis, training weights, bioinformatic data, the SPARROW implementation, and the Google Colab notebook are linked from this manuscript’s main GitHub directory: https://github.com/holehouse-lab/supportingdata/tree/master/2023/ALBATROSS_2023

## RESULTS

Our approach in developing ALBATROSS was to perform coarse-grained simulations of a set of training sequences that would enable an LSTM-BRNN model to learn the mapping between IDR sequence and global conformational behavior. To this end, four distinct phases in this process were required: (1) Selecting an appropriate force field, (2) Designing a library of synthetic sequences, (3) Performing simulations of those sequences, and 4) Optimizing deep learning models for sequence-to-ensemble mapping.

### The Mpipi-GG force field accurately recapitulates IDR ensemble dimensions

Before training a deep learning model for ensemble properties, we collected training data via coarse-grained simulations performed in the Mpipi-GG force field, a fine-tuned version of the previously published Mpipi forcefield^49^. We (and others) have had great success in using Mpipi to provide molecular insight into a range of systems^73, 74^. While Mpipi generally shows very good accuracy when compared with experiments, in performing initial calibration simulations, we noticed a few minor discrepancies between known experimental trends and Mpipi behavior (**Fig. S1-S5**). We made several small modifications to the underlying parameters, yielding a version of Mpipi we refer to as Mpipi-GG (see *Supplementary Information* for more details on force field fine-tuning).

To assess the accuracy of Mpipi-GG, we curated a set of 137 radii of gyration from previously published SAXS experiments on disordered proteins (**Fig. 1C**). Comparing the predictive power of Mpipi-GG to the original Mpipi force field for these sequences reveals comparable accuracy, with Mpipi-GG performing modestly better with an R^2^ of 0.921 vs. 0.896 for Mpipi, although both models are highly accurate (**Fig. 1C, Fig. S1C,D**). Given this accuracy we reasoned that we could leverage Mpipi-GG simulations to generate training data for deep-learning-based models to map IDR sequence chemistry to ensemble properties.

### Designing libraries of synthetic and biological IDRs for broad coverage of chemical space

Prior to performing simulations, we constructed a disordered protein sequence library with diverse sequence chemistries. We reasoned that a systematic exploration of IDR sequence space would enable our deep learning models to learn the complex underlying sequence-encoded conformational biases of disordered proteins. To build this initial synthetic library, we used GOOSE, our recently developed computational package for synthetic IDR design, to titrate across a range of sequence features known to impact IDR conformational behavior (**Fig 1D**, see *Methods*). Moreover, we opted to take advantage of GOOSE’s ability to focus compositional exploration to sequences predicted to be disordered, such that our initial library is centered on sequences predicted with high confidence to be IDRs.

Using GOOSE, we designed a library of synthetic sequences that systematically explore IDR hydropathy, overall charge, net charge, charge patterning (quantified by kappa (𝜅)), and the overall fraction of different amino acids (**Fig. 1D**, **Fig S6**). Lastly, we also added disordered sequences by specifying random sequence fractions. Collectively, we designed a library of 23,127 synthetic sequences across a broad sequence landscape. In addition to these synthetic IDRs, we also randomly selected 19,075 IDRs from common model system proteomes. In total, we collected a library of 42,202 disordered protein sequences. This training library covered broad chemical space in terms of the fraction of aliphatic and polar residues (**Fig. S6**) as well as the fraction of positively charged residues and aromatic residues (**Fig. S6**). An overview of the amino acid compositions and overall sequence complexity for the training library is summarized in **Fig. S6**. Moreover, we also ensured our sequence library had broad coverage of the sequence charge decoration (SCD) parameter defined by Sawle and Ghosh as well as the sequence hydropathy decoration parameters (SHD) (**Fig. S7**)^11, 75^.

### Training and evaluating an IDR sequence-to-ensemble deep learning model

After designing our training library of IDR sequences and both selecting and tuning our force field, we performed molecular dynamics simulations of all 42,202 sequences and calculated ensemble-average parameters of interest. Specifically, we focused on the radius of gyration, end-to-end distance, asphericity, and the scaling exponent and prefactor for the polymer scaling law fit the internal scaling data^23, 76, 77^. These data served as the foundation for training bidirectional recurrent neural networks with LSTM cells with PARROT for sequence-dependent property prediction tasks (see *Methods*). The collective group of these networks we term ALBATROSS.

We first began training the ALBATROSS R_g_ network. We leveraged the PARROT framework to train LSTM-based deep learning models on the simulated radius of gyration data. After optimizing hyperparameters (see Methods), we first began by confirming the ALBATROSS radii of gyration match the Mpipi-GG radii of gyration for the same experimental data presented in **Fig. 1C**. Indeed, we see strong quantitative agreement between ALBATROSS-derived radii of gyration and the experimental radii of gyration, despite the fact none of these sequences were in our training data (R^2^ = 0.998, **Fig. S8**). Inspired by this result, we next sought to assess how accurately ALBATROSS was able to predict the simulated Mpipi-GG R_g_ values on data unseen during training. Promisingly, when we evaluated our model on a held-out test set consisting of both synthetic and biological IDRs, we saw strong correlations in all cases (R^2^ = 0.995, **Fig. 2A**). Finally, ALBATROSS was comparable or more accurate than current state-of-the-art methods for radii of gyration prediction but enables a much higher throughput (thousands of sequences/second) than comparable models (**Fig. S8**). We next turned to evaluate the accuracy of our networks on the R_e_ prediction task. Similarly to the ALBATROSS R_g_ network, we observed a strong correlation between the ALBATROSS R_e_ and the Mpipi-GG R_e_ on the held out test set (R^2^ = 0.986, **Fig. 2B**).

**Figure 2.**
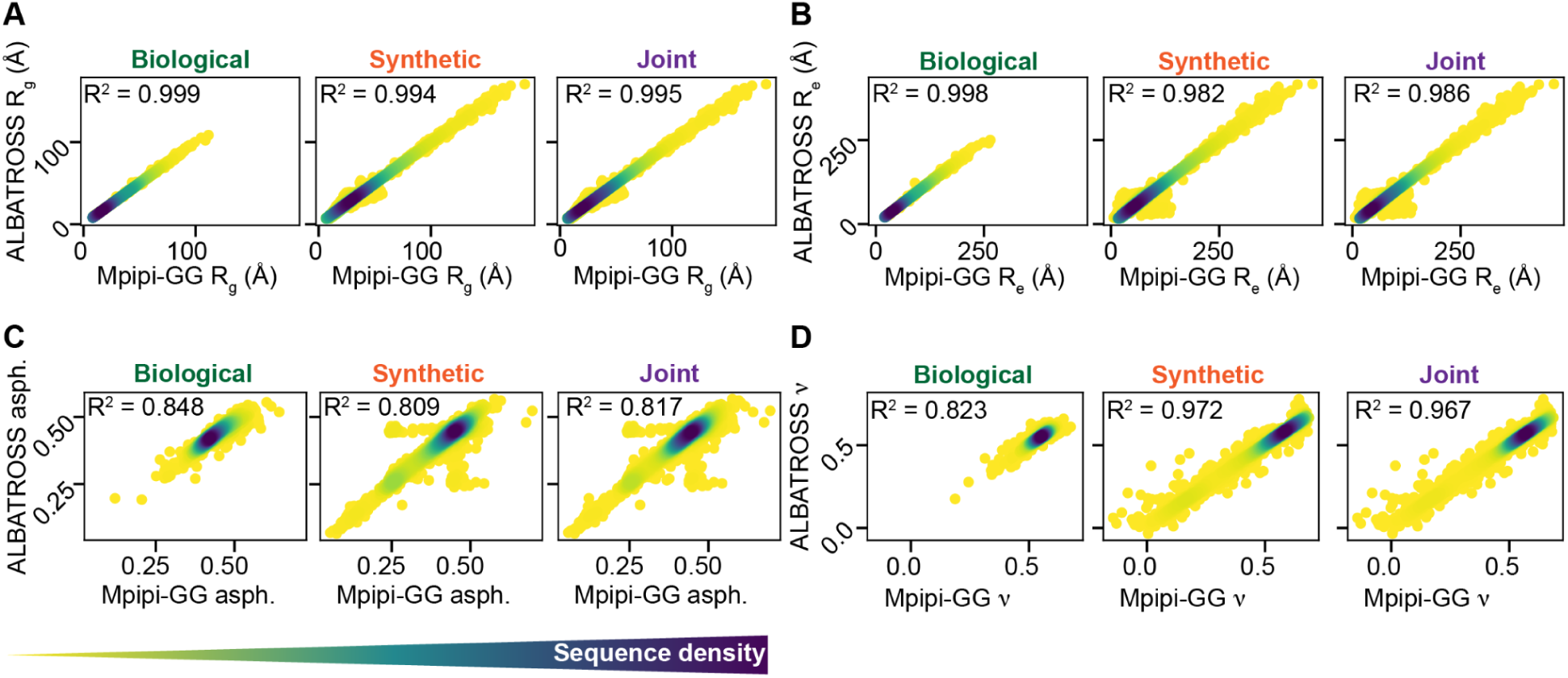
Assessing the ALBATROSS network accuracy on an independent test of both synthetic sequences and naturally occurring biological sequences. A, B, C, D) Accuracy of ALBATROSS in predicting R_g_, R_e_, Asphericity, and the polymer scaling exponent for previously unseen biological (left), synthetic (middle), and combined (right) sequences. For each correlation plot, a Gaussian kernel density estimation is used, where darker colors indicate regions where there are many sequences sharing a particular prediction value.

In addition to these R_g_ and R_e_ networks, we also trained networks for the mean asphericity, which displayed quantitative agreement on the test set (R^2^ = 0.817, **Fig. 2C**). Lastly, we trained predictors based on the two parameters obtained by fitting the internal scaling of the beads to a polymer scaling model; the scaling exponent and prefactor. The accuracy of the predictions from these networks was 0.967 and 0.930, respectively, on the independent set of test sequences (**Fig. 2D, Fig. S9**). In summary, the ALBATROSS networks perform well on both synthetically designed and naturally occurring IDRs suggesting our networks have learned the role of sequence chemistry for tuning IDR ensemble dimensions (**Fig. 2, Fig. S9**).

### ALBATROSS performance

While our four main networks are highly accurate, another benefit they provide is throughput for the systematic exploration of sequence-ensemble relationships. While coarse-grained simulations can take minutes, hours, or even days, ALBATROSS enables thousands of predictions per minute. A summary of our performance benchmarks on modest commodity CPU hardware is provided in **Fig. S10**, a criterion we focussed on, given many researchers do not have access to high-end GPUs. However, we note that one can compute R_g_ predictions for the entire human proteome in ∼8 seconds via our Google Colab notebook running on GPUs. As such, ALBATROSS offers an accurate and high-performance route to map sequence-ensemble relationships for R_e_, R_g,_ asphericity, and polymer scaling exponent and prefactor.

### Systematic assessment of sequence-to-ensemble properties

We next used ALBATROSS to assess how IDR sequence features influence global dimensions. Using GOOSE, we designed libraries of synthetic disordered sequences that systematically vary one sequence feature while holding others fixed. This strategy enables us to isolate and assess the average contribution of different sequence features; each data point on the panels in **Figure 3** reflects the average ensemble dimensions obtained from 100 distinct sequences with the same overall sequence features.

**Figure 3.**
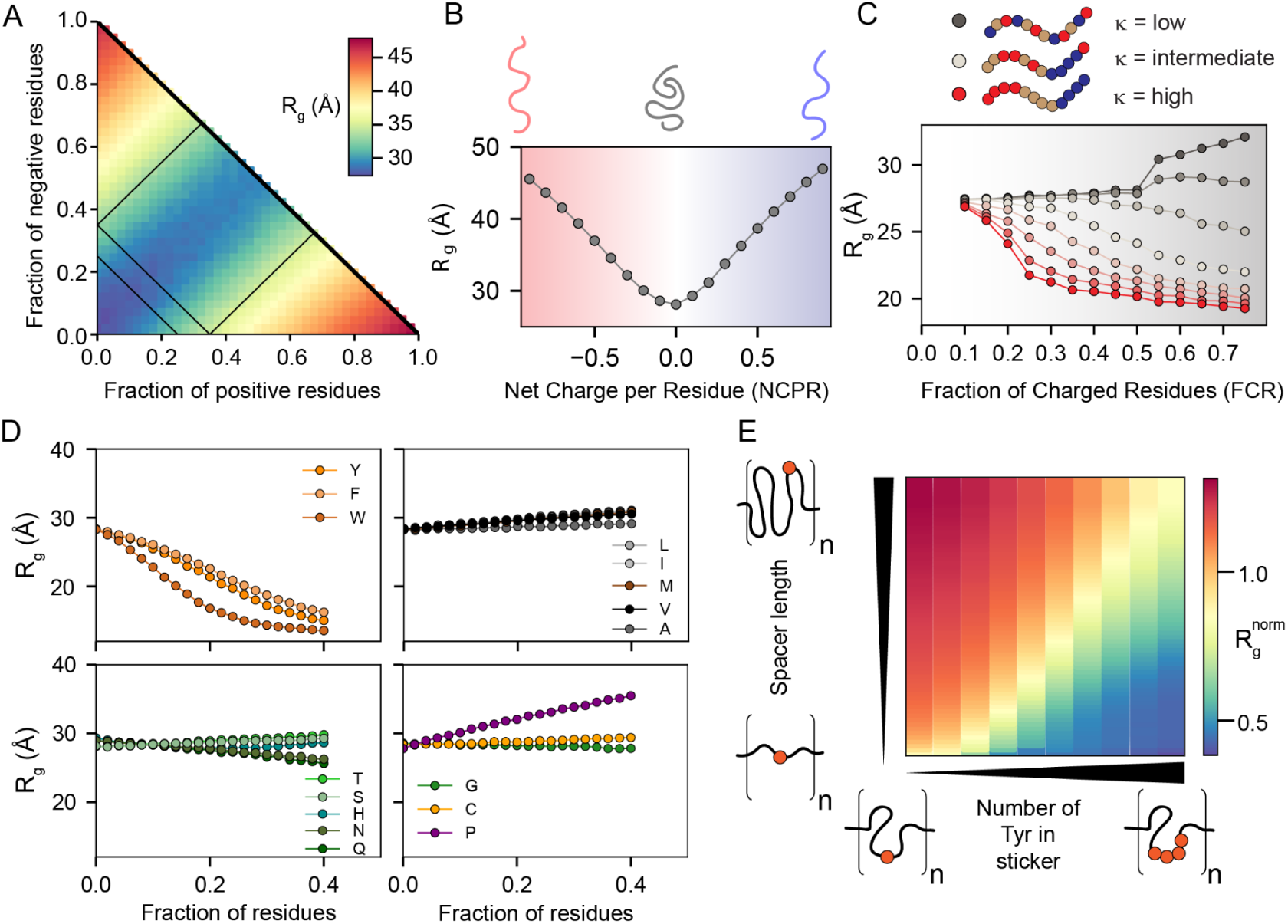
Sequence composition modulates the conformational preferences in disordered proteins. For panels A-D, each data point reports the average of many 100-residue synthetic disordered sequences with the specified composition. **A)** Diagram of states for weak to strong polyampholytes. Sequences are colored by a blue-to-yellow-to-red gradient based on their ALBATROSS radii of gyration. B) ALBATROSS radii of gyration as a function of net charge per residue. Both net negative (red) and net positive (blue) charged polyampholytes can drive chain expansion. **C)** The patterning of positively or negatively charged residues dictates the radius of gyration for highly-charged sequences but not those with a low fraction of charged residues. **D)** ALBATROSS radii of gyration as a function of the fraction of amino acid content for sixteen of the different amino acids. Aromatic residues drive compaction, while proline drives expansion. In each case, the fraction of other residues was held approximately fixed while one specific residue was systematically varied. **E)** Dependence of the normalized radius of gyration for sticker-spacer IDRs in which spacers are glycine serine repeats, and stickers are one or more tyrosine residues. The normalized radius of gyration is calculated as the ALBATROSS R_g_ divided by the R_g_ expected for a sequence-matched version of the protein behaving as a Gaussian chain (the AFRC model)^68^. Each sequence here contains 8 sticker-spacer repeats. Each repeat contains spacer regions (glycine-serine dipeptide repeats) that vary in length from 2 to 120 residues and sticker regions (poly-tyrosine repeats) that vary in length from 0 tyrosines to 8 tyrosines.

This analysis recapitulates and confirms a wide variety of sequence-to-ensemble relationships reported by many groups through computational and experimental studies over the last decade. In particular, our work highlights the importance of net charge in determining IDR global dimensions (**Fig. 3A, 3B**) and illustrates the fact that charge patterning becomes an increasingly important determinant of IDR dimensions as the overall fraction of charged residues increases (**Fig. 3C**)^11, 12, 21, 22, 24^. A systematic titration of individual amino acid fractions confirms that aromatic residues drive chain compaction (with tryptophan the strongest of the three), that proline residues drive chain expansion, and that glutamine (more than any other polar amino acid) drives intramolecular interactions and compaction (**Fig. 3D**) ^20, 23, 35, 78–80^. Finally, these analyses would suggest that aliphatic hydrophobes have a modest impact on IDR dimensions, a result consistent with prior work, although we caution that our predictions likely underestimate the hydrophobic effect (see Discussion) (**Fig. 3D**) ^81–83^. In summary, our conclusions here are largely concordant with prior work but generalize those conclusions from individual proteins or systems to the sequence-average properties.

In addition to titrating the aromatic fraction, we designed synthetic repeat proteins consisting of glycine-serine-repeat “spacers” and poly-tyrosine “stickers” ^84–86^. These synthetic IDRs allow us to assess how spacer length and sticker strength (tuned by the number of tyrosine residues in a sticker) influence chain dimensions. Our results demonstrate that both spacer length and sticker strength can synergistically influence IDR global dimensions (**Fig. 3E**). The dependence of the individual chain R_g_ on spacer length (y-axis) and sticker strength (x-axis) mirrors conclusions drawn from sticker-spacer architecture polymers from simulations and experiments ^23, 87–89^.

### Predicting emergent biophysical properties throughout the human proteome

Given ALBATROSS’ accuracy and throughput, we next performed large-scale bioinformatic characterization of the biophysical properties of disordered regions across the human proteome (**Fig. 4A, B**). Focusing on IDRs between 35 and 750 residues in length, we calculated normalized radii of gyration (**Fig. 4C**), normalized end-to-end distance (**Fig. 4D**), and asphericity (**Fig. 4E**). Normalization here was essential to account for the variability in absolute radii of gyration with sequence length, and was achieved by dividing the ALBATROSS R_g_ with the sequence-specific R_g_ expected if the IDR behaved as a Gaussian chain^68^. These analyses suggest that most IDRs behave as relatively expanded chains, although we recognize there are likely several important caveats to this interpretation (see *Discussion*). Assessing the absolute radius of gyration vs. IDR length, the majority of more compact IDRs are enriched for aromatic residues (**Fig. 4F**). Indeed, plotting the asphericity (a measure of IDR ensemble shape) vs. the normalized radius of gyration and coloring by either the fraction of aromatic residues (**Fig. 4G**) or the absolute net charge and the fraction of proline residues (**Fig. 4H**) suggest that IDRs with an ensemble that is expanded and elongated have a net charge and/or are enriched for proline, whereas IDRs with an ensemble that is compact and more spherical are enriched for aromatic residues. Segregating IDRs into the 1000 most compact and 1000 most expanded sequences reveals that compact IDRs tend to be depleted in proline residues and have a low NCPR, whereas those that are expanded are enriched in proline and/or have an absolute NCPR, although we found many examples of proline-rich charge depleted IDRs that were relatively expanded. Taken together, our analysis of the human IDR-ome mirrors insights gleaned from the analysis of synthetic sequences in **Fig. 3**.

**Figure 4.**
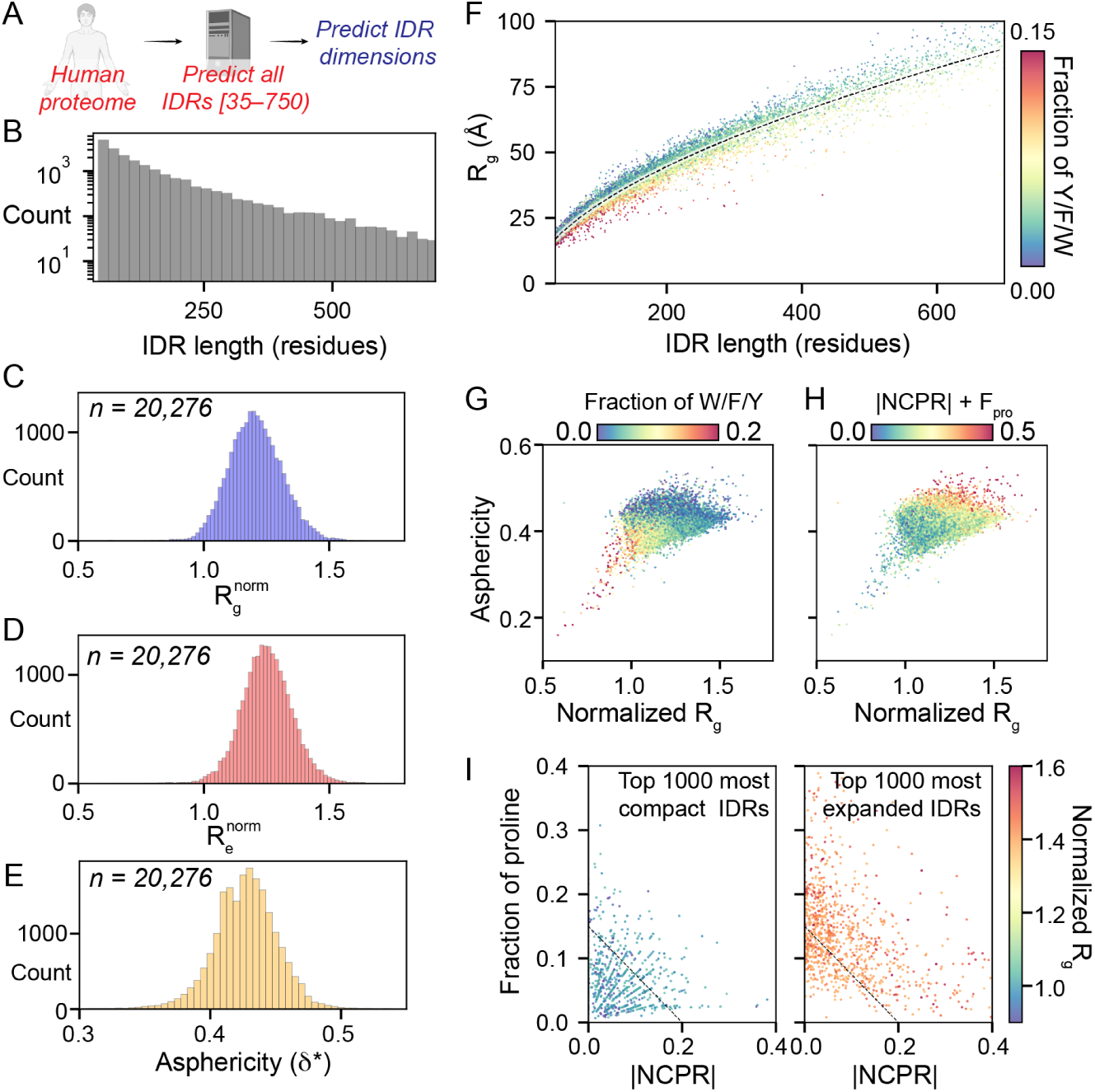
Human proteome-wide biophysical characterization of predicted IDRs. A) ALBATROSS was used to perform sequence-dependent ensemble predictions for all IDRs in the human proteome. **B)** Histogram of all human IDRs ranging from 35 to 750 residues. **C)** Normalized mean ALBATROSS R_g_ distribution for all human IDRs. **D)** Normalized mean ALBATROSS R_e_ distribution for all human IDRs. **E)** Mean ALBATROSS asphericity distribution for all IDRs in the human proteome. **F)** Mean ALBATROSS radius of gyration as a function sequence length. Individual data points are colored by the fraction of aromatic residues in the sequence. The dashed line represents the fitted scaling law, which reports an apparent scaling exponent of 0.56. Deviations above and below this line suggest sequence-specific expansion or compaction, respectively. **G)** Full distribution of human IDRs plotted in terms of the normalized radius of gyration and asphericity, colored by the fraction of aromatic residues. **H)** Full distribution of human IDRs plotted in terms of the normalized radius of gyration and asphericity, colored by the absolute net charge per residue plus the fraction of proline residues. **I)** Top 1000 most compact (left) and top 1000 most expanded (right) IDRs plotted in terms of the fraction of proline residues and absolute net charge per residue.

### Characterizing local dimensions of IDR subsequences

Our proteome-wide analysis in **Fig. 4** focused on ensemble-average properties calculated for entire IDRs. While convenient for revealing gross properties, we reasoned that for large (200+ residue) IDRs, it may be more informative to assess local conformational behavior with a sliding-window analysis. To this end, using a window size of 51 residues, we calculated the local end-to-end distance across every 51-mer fragment in the human proteome, enabling us to extract the 2,146,400 51-mer fragments that lay entirely within every IDR (**Fig. 5A**). The distribution of normalized end-to-end distances is tighter than the corresponding distribution for full-length IDRs, with 195,555 (i.e., around 10%) of subregions behaving as polymers more compact than a corresponding Gaussian chain (normalized R_g_ < 1) (**Fig. 5B**).

**Figure 5.**
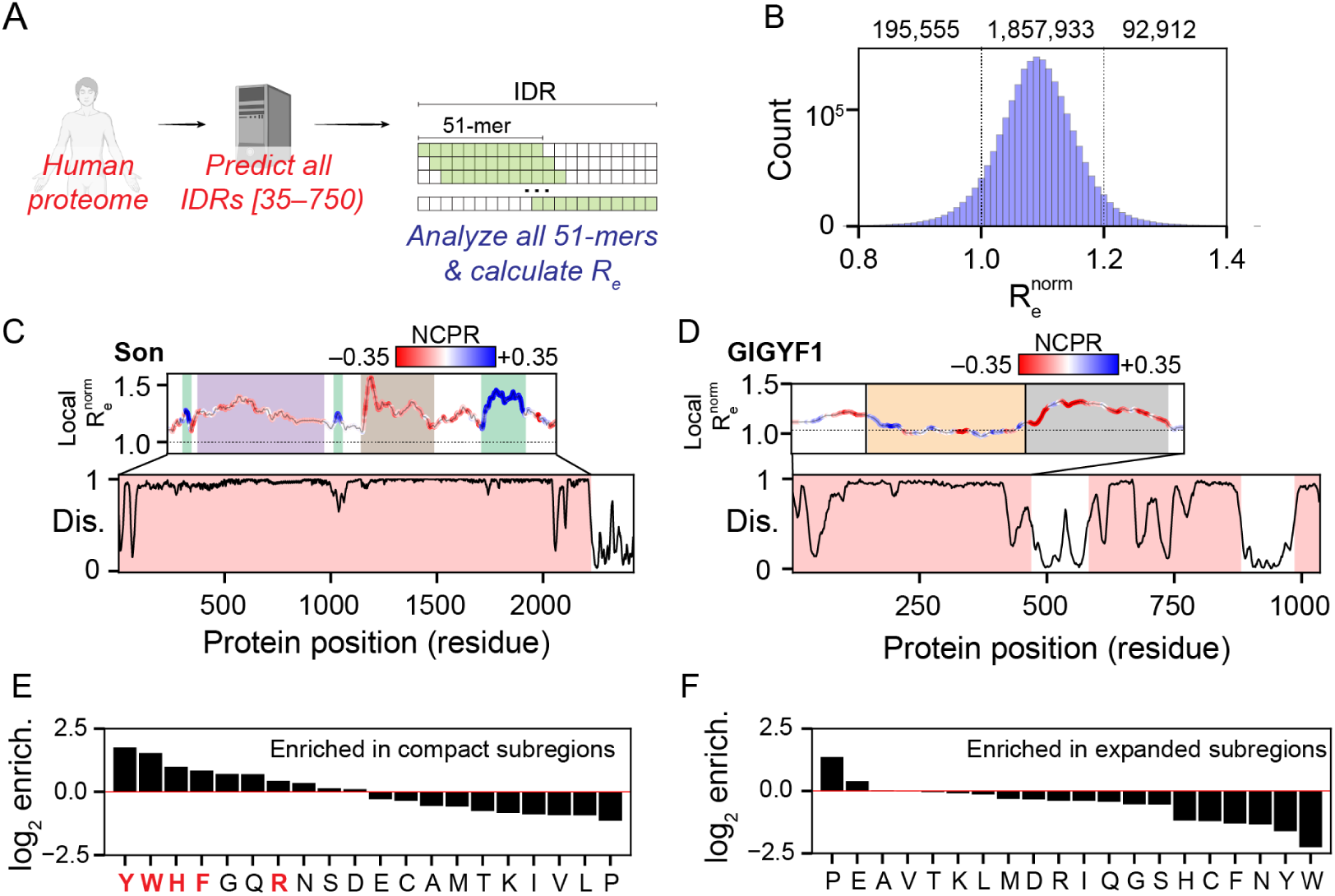
Local analysis of disordered protein subregions reveals sequence-dependent expansion and compaction. A) Graphical summary illustrating the sliding window subregion analysis presented in this figure. **B)** Distribution of the normalized end-to-end distance obtained from all 51-residue subfragments within IDRs in the human proteome. **C)** Linear analysis of local subregions in the 2227 residue IDR from the nuclear speckle protein Son, with conformationally-distinct subregions highlighted (UniProt: P18583). **D)** Linear analysis of local subregions in the 469-residue IDR from the cytosolic GIGYF1, with conformationally-distinct subregions highlighted (UniProt: O75420). **E)** Log_2_-fold enrichment for amino acids found in compact subregions. Residues implicated in RNA binding are highlighted in red. **D)** Log_2_-fold enrichment for amino acids found in expanded subregions.

The linear assessment of local dimensions enables the demarcation of conformationally-distinct subdomains within an IDR. As a proof-of-concept, we plotted the normalized local end-to-end distance for two large IDRs, revealing distinct subregions within each (**Fig. 5C, D**). First, we analyzed the 2227 residue IDR from the nuclear speckle protein Son, identifying distinct subregions with specific conformational properties that map to previously analyzed subregions within the sequence (**Fig. 5C**)^90^. Second, we analyzed the N-terminal IDR of GIGYF1, a highly disordered protein with a potential role in Type II diabetes ^91–93^. The N-terminal IDR in GIGYF1 contains three subregions, an expanded N-terminal region that may fold upon binding (residues 1-90), a comparatively compact central region (residues 91-280), and an expanded C-terminal acidic region (residues 281-469). The ability to – from sequence alone – demark potential subdomains within an IDR paves the way for more sophisticated mutagenesis studies, as well as the ability to predict if and how mutations might influence local conformational behavior, and potentially, molecular function.

Finally, we used the set of ∼2 million IDR subregions to assess which residues were enriched in expanded or compact IDRs (**Fig. 5E, D**). Enrichment was assessed based on the fraction of the twenty amino acids in subregions taken from the top/bottom 2.5% of all subregions with respect to normalized end-to-end distance, compared to the overall fraction for all subregions. Aromatic residues, histidine, arginine, glycine, and glutamine were all found to be enriched in compact subregions. In contrast, proline and glutamic acid were found to be enriched for expanded subregions. Intriguingly, several of the residues most strongly enriched for compact IDRs match those residues known to engage in RNA binding^73, 94–97^. Moreover, a gene ontology analysis for proteins with 10 or more compact subfragments found strong enrichment for RNA binding (**Table S1**). In contrast, we saw no obvious patterns in proteins that possessed expanded subregions (**Table S2**). Taken together, our analysis suggests IDRs that favor intramolecular interaction may share a common molecular function in RNA binding, whereas those that are highly expanded likely play a variety of context-specific roles.

### High-throughput IDR ensemble informatics across yeast evolution

Having demonstrated the accuracy and throughput of ALBATROSS in conducting broad, proteome-wide analyses, we next sought to showcase ALBATROSS’ unique advantages for structural bioinformatics of disordered proteins. In particular, we were motivated by recent work demonstrating that IDRs can conserve global dimensions in spite of variation in amino acid sequence, as reported for a linker region in the viral protein E1A^72^. Compelled by this specific example, we suspected that there would be other instances in nature in which functionally-important IDR dimensions are evolutionarily conserved across divergent homologs.

To test this, we analyzed evolutionarily-related IDRs across a wide-ranging set of yeast species. Using the *S. cerevisiae* proteome as a reference, we aligned and extracted 2302 sets of homologous IDRs >40 residues from 20 yeast proteomes, totaling 49,335 IDRs (**Fig. 6A**). We predicted the R_e_ for all IDRs, and used the standard deviation of these predicted R_e_ values to quantify the conservation of IDR dimensions. We quantified homolog sequence divergence using two approaches: by computing the variation of IDR sequence lengths and by scoring the sequence similarities from the multiple sequence alignment (see *Methods*). In line with what has been reported elsewhere, both of these approaches for scoring sequence similarity show that homologous IDRs in our dataset are significantly more divergent than homologous folded domains (Mann-Whitney U test, p < 0.001) (**Fig. S11**) ^70, 98–100^.

Looking at all of the sets of homologous IDRs, we see a clear relationship between sequence similarity and R_e_ conservation (**Fig. 6B, S12**). As expected, IDRs with more divergent sequences tend to possess larger variations in R_e_, however there are a number of homologs that exhibit tightly coupled R_e_ relative to their sequence (dis)similarity, even compared to E1A^72^ (**Fig. 6B, S12**). While characterizing all of these homologs is outside of the scope of this work, we did perform a deeper analysis into two of these candidate IDRs in order to validate this observation.

**Figure 6.**
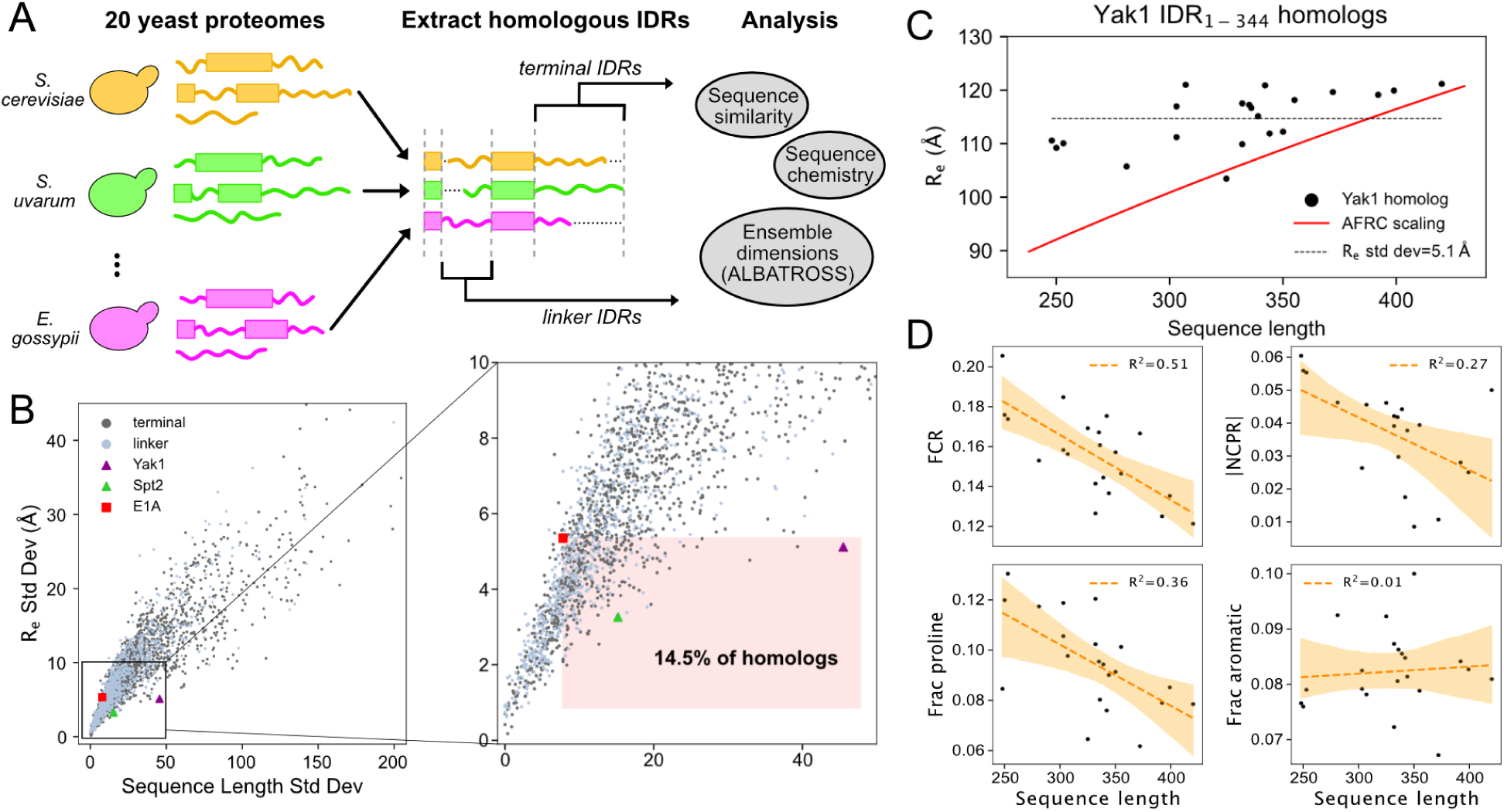
Applying ALBATROSS on homologous yeast IDRs identifies instances where primary sequences diverge yet IDR dimensions are maintained. A) Graphical summary illustrating how sets of homologous proteins from 20 yeast proteomes were aligned, then IDRs were extracted and analyzed using ALBATROSS and bioinformatics. B) For each set of homologous IDRs, standard deviation of the predicted R_e_ values is plotted with the standard deviation of sequence length (in number of amino acids). Linker IDR homologs and terminal IDR homologs are plotted in blue and gray, and several specific IDR homologs are marked. The right panel shows a zoomed-in inset, with a region denoting homologs with more divergent sequences and less variable dimensions than E1A are highlighted. C) Sequence length and ALBATROSS-predicted R_e_ for Yak1 and its yeast homologs. The gray dashed line denotes the mean R_e_. The red line denotes R_e_ as a function of sequence length for an Analytical Flory Random Coil polymer scaling model. D) Yak1 homolog sequence length compared to various protein sequence features. Line of best fit and 95% CI denoted in orange. FCR=fraction of charged residues; |NCPR|=absolute value of net charge per residue.

The first system we looked at was the N-terminal IDR of the DYRK-family kinase Yak1 and its homologs. Despite large variation in sequence length (248 - 420 residues), the Yak1 homologs displayed strikingly conserved dimensions, with all but two sequences having predicted R_e_ between 109 and 122 Å (**Fig. 6C**). In contrast, basic polymer models predict R_e_ differences of >27 Å across this range of sequence lengths, suggesting that there may be evolutionary constraints on the R_e_ of the N-terminal Yak1 IDR. Analysis of the IDR sequences reveals several trends in sequence features that may explain the buffering of chain dimensions. As sequence length increases, there is a decrease in the fraction of proline and charged residues and a modest trend towards a neutral net charge (**Fig. 6D, Table S3**). Each of these features are associated with chain compaction ^72^(**Fig. 3**, **Fig. 4**, **Fig. 5**). Other trends, such as a modest increase in 𝜅, and changing sequence composition of polar and aliphatic residues may also modulate the dimensions and properties of the Yak1 homologs (**Fig. S13**).

We also examined the disordered linker (residues 57-373) from Spt2, a histone chaperone associated with chromatin remodeling during transcription. Like Yak1, the Spt2 IDR homologs have relatively constrained R_e_ despite spanning 186 to 239 residues in length (**Fig. 6B, S12, S14**). Through analysis of the sequence features of the Spt2 homologs, their dimension of the longer sequences appear to be modulated by an increase in proline and aliphatic content, and a more neutral net charge. Additionally, all of the homologs have high κ values which may buffer the overall chain dimensions (**Fig. S14, Table S4**).

## DISCUSSION

Intrinsically disordered proteins and protein regions (IDRs) are ubiquitous, yet the absence of a fixed 3D structure coupled with limited sequence conservation has challenged conventional routes for mapping between protein sequence and molecular function. Given IDR function can be influenced or even dictated by the sequence-encoded conformational biases, a robust understanding of sequence-ensemble relationships remains an important feature for interpreting how IDRs conduct their cellular roles ^19, 72, 101^.

Here, we present ALBATROSS, a deep learning approach trained on coarse-grained simulations that allow for direct prediction of ensemble-average global dimensions from protein sequences. While there are several caveats that should be considered, ALBATROSS enables us to assess sequence-to-ensemble relationships for both synthetic and natural IDRs. By providing ALBATROSS as both a locally-installable Python package and an easy-to-use Google Colab notebook, we aim to lower the barrier for sequence-to-ensemble predictions for single IDRs or for entire proteomes.

Our proteome-wide analysis suggests that IDR expansion can be driven by net charge, proline residues, or a combination of the two (**Fig. 4I**). In contrast, the subset of amino acids (Y/W/F/H/R/G/Q) enriched in compact IDR subregions overlap strongly with those residues previously reported to engage in RNA binding (**Fig. 5E**). Previous work has shown that disordered regions can chaperone RNA, both in isolation and in the context of biomolecular condensates^73, 102–105^. Interestingly, these same RNA binding residues are also over-represented in IDR subregions that can drive phase separation *in vitro* and form condensates *in vivo* ^23, 78, 85, 106–110^. One interpretation of these observations is that compact IDRs have evolved to self-assemble and recruit RNA into condensates. Another interpretation is that these RNA-binding IDRs are constitutively bound to RNA in cells where they exchange compaction-driving intramolecular protein:protein interactions for expansion-driving intermolecular protein:RNA interactions. Under this interpretation, compact IDRs are only compact in an unphysiological RNA-free context, such that they expand to envelop and chaperone RNA molecules while themselves being reciprocally chaperoned by RNA. These interpretations are not mutually exclusive, nor do they prohibit a model in which RNA chaperoning requires many copies of RNA-binding proteins forming dynamic condensates.

Our analysis of yeast homologs highlights two specific cases where IDR dimensions appear to be conserved across evolution despite substantial divergence in primary sequence, consistent with previous studies^72^. The homologs of the Yak1 N-terminal IDR homologs have more constrained R_e_ than we would expect based on polymer models (**Fig. 6C**). In the literature, Yak1 kinase activity has been shown to be regulated by its N-terminal IDR, through both intra- and intermolecular interactions^111^. We hypothesize that maintaining a narrow range of end-to-end dimensions of the N-terminal IDR across homologs could be important for preserving autophosphorylation capabilities and for facilitating specific, multivalent interactions with 14-3-3 proteins^111, 112^. The IDRs of the histone chaperone Spt2 also maintain similar R_e_ values across divergent homolog sequences (**Fig. S14**). Spt2 has shown to interact with histone proteins and a variety of chromatin remodelers, with its IDR potentially playing an important role in these binding events^113, 114^. Thus, we hypothesize that the Spt2 IDR dimensions are functionally conserved across homologs in order to preserve the protein’s ability to act as a multivalent molecular scaffold. Beyond Yak1 and Spt2, our analyses suggest there are likely other IDRs with evolutionary constraints on their dimensions. Although we need further experiments to validate any of these claims we make regarding mechanism, we believe that this analysis illustrates ALBATROSS’s utility in generating hypotheses and identifying candidates in large biological datasets.

Recent work from several groups touches on ideas or results that dovetail well with our own. As a proof-of-principle, Janson et al. trained a generative adversarial network to predict ensemble properties for coarse-grained simulations (idpGAN) ^115^. This study also demonstrated the potential for multi-resolution models that interpolate between coarse-grained and atomistic simulations. Leveraging and interpolating between these multiscale models to generate thermodynamically distinct but realistic conformations may be a promising approach to enhance conformational sampling^116, 117^. In parallel, Chao *et al.* presented a novel approach to represent IDR ensembles and train several different machine learning architectures to predict global dimensions from sequence^118^. This work suggests that representing IDRs in a length-free way using the Bag of Amino Acids representation may be useful to capture sequence-specific effects and offers some advantages in model training. Finally, Tesei & Trolle et al. recently performed an analogous assessment of the human IDR-ome using the CALVADOS2 force field^48, 53, 119^. Despite using a different force field, the correlation between CALVADOS2 simulations of the human proteome and ALBATROSS predictions is high, with root mean squared errors within the range of experimental error (**Fig. S15**, R^2^ = 0.98, RMSE = 3.68 Å, n=29,998, ALBATROSS prediction time for all IDRs ∼200 seconds on a CPU). Moreover, we arrive at similar conclusions for the propensity for relatively expanded IDRs, the importance of net charge, charge patterning, and aromatic residues in tuning overall dimensions, and the association between RNA binding proteins and compact IDRs. Overall, the distribution of IDR dimensions from CALVADOS2 is slightly more compact than from Mpipi-GG, a difference we suspect reflects an underestimation of aliphatic residue interactions in the Mpipi-GG force field. Nevertheless, the general trends between the two studies show good agreement, a compelling result given the differences in approaches, force fields, and assumptions.

While our benchmarks demonstrate the predictive power of simulations performed using Mpipi-GG and of ALBATROSS, there are a few important limitations to consider. Mpipi-GG is a one-bead-per-residue, coarse-grained force field that assumes an isotropic interaction potential. Despite this simplifying assumption, many independent studies have confirmed that coarse-grained models are able to capture global ensemble properties of IDRs with reasonably good accuracy^48, 49, 51, 53, 120^. Nevertheless, we suggest a few specific caveats that should be considered when evaluating Mpipi-GG simulations or ALBATROSS predictions. Firstly, we likely underestimate the impact of solvation effects on charged amino acids, such that highly-charged, net-neutral IDRs are likely more compact than they should be. Secondly, our coarse-grained model and predictors do not account for transient secondary structure elements, a pervasive source of local conformational heterogeneity in many IDRs. Finally, we likely underestimate the hydrophobic effect for aliphatic residues, an intrinsically challenging phenomenon to capture in coarse-grained force fields for IDR simulations. These two final points mean we likely overestimate the predicted dimensions of IDRs that possess hydrophobicity-driven secondary structure, a caveat that should be carefully considered for IDRs enriched for helicity-promoting and/or aliphatic residues.

Our decision to focus on an LSTM-BRNN architecture for training was motivated by the desire to develop trained networks that were performant (10-50 sequences/second) on CPU commodity hardware. While more complex architectures (e.g., transformer-based networks) may offer more accurate predictors, we see two central limitations here. First, transformer-based architectures are memory intensive, and although some low-memory transformer-based architectures exist, most pre-trained biological transformers have memory requirements that scale quadratically with sequence length^121–126^. These large memory requirements can be prohibitive on commodity hardware, and we wanted to focus on developing and distributing portable tools for the community. Second, our LSTM-based architecture generates predictions that are already quite accurate. The error associated with our predictions is on the order of the experimental error (0 - 4 Å), so treating model architecture as a tunable hyperparameter for the performance of these prediction tasks, while an interesting question, did not merit further experimentation. Finally, combining an LSTM-based sequence-to-ensemble predictor with our high-throughout LSTM-based disorder predictor (metapredict V2-FF) ensures parity in disorder prediction and ensemble prediction. With this in mind, our IDR-ome predictor notebook enables proteome-wide predictions in minutes, democratizing high-throughput unstructural bioinformatics.

## CONCLUSION

In this work, we present ALBATROSS, an accessible and accurate route to predict IDR global dimensions from sequence. Our results are in good agreement with prior experimental and recent analogous computational work, suggesting that ALBATROSS offers a convenient route to obtain biophysical insight into IDR sequence-ensemble relationships.

## ACKNOWLEDGEMENTS

We thank members of the Holehouse lab for extensive discussions on many aspects of this work. We thank Shubh Minhas for the ALBATROSS logo. We are also indebted to Kresten Lindorff-Larsen for his willingness to delay preprinting of their manuscript such that both preprints could reference one another. We also thank past and present members of the Pappu lab, Sukenik lab, Soranno lab, and Mittag lab for many illuminating discussions on sequence-ensemble relationships over the years. Funding for this work was provided by the Human Frontiers in Science Program (HFSP RGP0015/2022) to A.S.H. by the Longer Life Foundation, an RGA/Washington University in St. Louis Collaboration to A.S.H., and by the National Science Foundation with award 2128068 to A.S.H. JML was supported by the National Science Foundation via grant number DGE-2139839. DG was supported by the NSF via grant number DGE-2139839. We also thank members of the Water and Life Interface Institute (WALII), supported by NSF DBI grant #2213983, for helpful discussions.

