## Supplementary Information for "Direct Prediction of Intrinsically Disordered Protein Conformational Properties From Sequence: Version 3 [2023-05-28]"

#### 1. Extended methods

##### *Mpipi fine-tuning*

Mpipi is a one-bead-per-residue coarse-grained force field that was parameterized via a bottom-up, data-driven approach using statistics obtained from the PDB coupled with quantum mechanical calculations and all-atom simulations to derive parameters for a Wang-Frenkel (WF) potential **(Equation 1.1-1.2)** [1,2].

$$\phi(r) = \epsilon\,\alpha\left(\left[\frac{\sigma}{r}\right]^{2\mu} - 1\right)\left(\left[\frac{R}{r}\right]^{2\mu} - 1\right)^{(2\nu)} \qquad \textbf{Equation 1.1}$$

where

$$\alpha = 2\nu\left(\frac{R}{\sigma}\right)^{2\mu} \times \left(\frac{1+2\nu}{2\nu\left[\left(\frac{R}{\sigma}\right)^{2\mu} - 1\right]}\right)^{(2\nu+1)} \qquad \textbf{Equation 1.2}$$

Where $R = 3\sigma$ and $\nu = 1$, as defined in the original Mpipi paper.

The WF potential provides a computationally convenient (efficient to compute, intercepts with 0 at long intermolecular distances) closed-form alternative to the more commonly used Lennard-Jones potential. Given the data-driven approach used for parameterization, Mpipi benefits from explicitly encoded inter-residue interaction values (i.e., $\varepsilon_{i,j}$ values) for all unique pairs of amino acids. This is in contrast to most other one-bead-per-residue force fields, where intrinsic residue-specific interaction strengths (i.e., $\varepsilon_i$ or $\lambda_i$) are defined, and inter-residue interaction energies are then computed via so-called 'mixing rules'. While Mpipi offers improved flexibility for capturing chemically complex interactions, the model also has many more parameters than most conventional force fields (i.e., $[n^2 + n]/2$ interaction parameters for a

model with *n* amino acids). As such, despite the excellent accuracy of the original model, we sought to determine if Mpipi could be further improved for certain sequence chemistries.
We focused on four specific groups of pairwise interactions to fine-tune Mpipi. In doing so, we developed an augmented version we refer to as Mpipi-GG. In particular, we strengthened Gly:Gly and Gly:Ser interactions, weakened aromatic:charge interactions, increased the excluded volume of proline residues, and reparameterized aliphatic residues to have increased hydrophobicity (**Fig. S1**).

The adjustment to proline was motivated by the observation that upon simulation with the original Mpipi parameters, many proline-rich IDRs were too compact compared with experiments (**Fig. S2A, B**). Proline predominantly drives IDR expansion via backbone restrictions and favorable solvation[3–6]. We reasoned that tuning the proline σ parameter (i.e., its excluded volume) would enhance its expansion-driving effects. To this end, after systematically titrating potential σ values and comparing the outcome of altering the parameters with all-atom simulations (**Fig. S2C**), we increased the proline σ by 33% for all pair-wise proline interactions, as shown in **Fig. S2B**. Applying this fix improved accuracy with respect to proline-rich IDRs with minimal loss of accuracy for other IDRs (**Fig. S2D**).

Following our adjustment of proline, we examined several polar-rich homo- or dipolymeric tracts for which experimental data have previously been obtained; poly-(GS), poly-(G), poly-(S), and poly-(Q). Previous work established that sufficiently long polyglutamine (poly-(Q)) tracts from compact globules, consistent with results from Mpipi[7]. However, we noticed that both poly-(GS) and poly-(G) scaled as a self-avoiding random walk ($\nu$ = ~0.60), despite the fact experimental work has suggested poly-(GS) behaves akin to a Gaussian chain ($\nu$ = ~0.5-0.55) and poly-(G) forms compact ensembles ($\nu$ = ~0.4) [8–11] (**Fig. S3A**). To address this discrepancy, we performed a titration series for poly-(G) chains. We tuned the G:G interactions by titrating the strength of the glycine-glycine attractive parameter in the WF potential ($\varepsilon_{G,G}$) for a poly-$(G)_{80}$ chain (**Fig. S3B**). Fitting these data to a coil-to-globule transition, we extracted the interaction strength (2.19x the original $\varepsilon_{G,G}$) that gave an apparent scaling exponent of 0.39, in line with previous experiments (**Fig. S3C**)[11,12]. Having established the correction factor for $\varepsilon_{G,G}$, we applied this same factor to the $\varepsilon_{G,S}$, such that poly-(GS) shows a slightly more compact scaling ($\nu$ = ~0.58) but is substantially more compact in terms of absolute dimensions, in better agreement with experiment (**Fig. 3D**). While this is not in perfect agreement with experimental work, it is an improvement on prior behavior. Without a reliable benchmark for poly-(S) we did not tune the $\varepsilon_{S,S}$ and instead focused on the $\varepsilon_{G,G}$ and $\varepsilon_{G,S}$ values. In summary, these changes provide a modest improvement in the expected polymer scaling behavior for glycine-rich sequences compared to the original Mpipi parameters.

To tune charge-aromatic interactions in Mpipi-GG we used aromatic-aromatic interactions as a benchmark. We compared the fraction of charged residues (FCR) and the fraction of aromatic residues with deviations in radii of gyration from experiment ($\Delta R_g$). Our analysis revealed that

Mpipi tended to over-compact sequences with greater aromatic and charge fractions (**Fig S4A**). We next plotted the pairwise WF-potentials, which suggested that the arginine, aspartic acid, and glutamic acid to aromatic interaction strengths were overestimated, likely driving this compaction (**Fig S4B**). Therefore, we tuned the $\varepsilon_{RED,FYW}$ values by systematically titrating to better fit the radii of gyration for these sequences (**Fig 4A, left**). The final $\varepsilon_{RED,FYW}$ value was 60% lower for the Mpipi-GG parameters. We confirmed our modifications better matched experimental radii of gyration by comparing the root mean squared error (RMSE) between simulated and experimental $R_g$ values at different fractions of aromatic residues for the Mpipi-GG and original parameters shown in **Fig. 4A** (**Fig S4C**).

The final set of parameter modifications focuses on aliphatic residues. Aliphatic residues in the original Mpipi force field have very weak interaction strengths. To incorporate hydrophobicity, we made use of the Kyte-Doolittle hydropathy ($KD_{hyro}$) scale to reparameterize pairwise aliphatic interactions, such that $\varepsilon_{ij}$ values are proportional to the sum of $KD_{hyro\ i+j}$ for a pairwise aliphatic interaction of i:j. **(Fig. S5A, B)**. Specifically, we modulated the aliphatic $\varepsilon_{AMLVI,AMLVI}$ values to strengthen aliphatic:aliphatic interactions **(Fig. S5B).**

In summary, small changes were made to parameters associated with twelve of the twenty natural amino acids: proline, glycine, arginine, aspartic acid, glutamic acid, phenylalanine, tyrosine, tryptophan, alanine, valine, isoleucine, leucine, and methionine. All changes made to the interaction matrix can be visualized in **Fig. S1**, which reports on the change in the overall interaction parameter between Mpipi-GG and Mpipi. The overall interaction parameter reports the net integral of both the short-range (Wang-Frenkel) and long-range (Coulombic) interaction potentials. We emphasize that these changes were made explicitly with single-chain behavior in mind and have not been tested in terms of their impact on phase behavior. As such, while the original Mpipi model may be preferable for studying two-phase systems, we proceeded to use Mpipi-GG for single-chain sequence-ensemble predictions.

***SAXS data used for comparison***

We assessed the accuracy of Mpipi using extant SAXS data obtained from the literature[13]. Wherever possible, we re-analyzed primary scattering data to ensure that reported values matched the radii of gyration reported in prior publications.

***IDR sequence library design***

To construct *bona fide* disordered protein sequences, we leveraged the software package GOOSE, which enabled us to construct libraries of rationally designed disordered proteins with specific yet broad sequence chemistries. Therefore, we first designed disordered sequences with varying Fractions of Charged Residues (FCR), Net Charge per Residue (NCPR), and Kyte-Doolittle hydropathy scale values. In addition, we also generated disordered sequences with randomly assigned but specific amino acid fractions (where remaining amino acids were unrestrained) to sample across the sequence space that is accessible to disordered regions.

Finally, we also ensured that we had broad coverage of charge distribution in our generated sequences by titrating across kappa, a charge asymmetry parameter where higher values mean greater charge asymmetry.

***All-atom Excluded Volume (EV) simulations***

Coarse-grained excluded volume (EV) simulations were performed in Mpipi-GG by adjusting the epsilon and sigma parameters such that the interaction potential overlaps for the repulsive component of the function but flattens to zero for distances greater than $\sigma$ (**Fig. S16**). All-atom EV simulations for polyproline (**Fig. S2**) were performed using the CAMPARI simulation engine (V2) https://campari.sourceforge.net/) and the ABSINTH implicit solvent model[14]. EV simulations were performed as done previously[3,15]. Briefly, EV simulations involve scaling the attractive Lennard-Jones component, the solvation component, and the electrostatic component of the ABSINTH Hamiltonian to zero, such that the only determinant of the underlying ensemble reflects the excluded volume dictated by the repulsive component of the Lennard-Jones potential.

**Scaled network training**

For both the radius of gyration and the end-to-end distance networks, we trained BRNN-LSTM networks with and without sequence length normalization. For the normalized variations, we performed normalization by taking the respective metric and dividing it by the square root of the sequence length. The radius of gyration (or analogously the end-to-end distance) for a polymer can be defined as $R = A_0 \times N^{\nu}$ where R is either the radius of gyration or end-to-end distance, N is the length of the sequence, and $\boldsymbol{\nu}$ is the scaling exponent. A Gaussian chain is a chain that scales with $\boldsymbol{\nu}$ as 0.5. Therefore, to obtain the length-independent (i.e., sequence chemistry) contribution to the chain dimensions, one can normalize R by the root of the sequence length to derive the following relationship $\frac{R}{\sqrt{(N)}} = A_0 \times N^{(\nu-0.5)}$. This scaling normalizes the measure in polymer space and standardizes the ensemble dimension such that length is a less dominant factor of the learned network. For each scaled network, we followed the same 5-fold cross validation procedure for hyperparameter tuning and the final network weights were selected from the lowest validation loss across 750 epochs.

**ALBATROSS distribution**

In addition to providing a locally installable implementation of ALBATROSS via SPARROW, we also created a point-and-click style interface for ALBATROSS hosted on Google Colab to eliminate the software barrier of entry for users. By leveraging the cloud computing resources provided in Google Colab, we enable the accurate prediction of IDR conformational properties from sequence from anywhere in the world with an internet connection - even a smart device. Moreover, our Google Colab implementation enables users to specify either a single sequence or upload a fasta file of disordered protein sequences for ALBATROSS predictions. This means users can leverage the unique throughput of ALBATROSS predictions without even needing to write code to construct complex bioinformatic pipelines. Additionally, all predictions are filterable by numerical ranges for that property. For example, if one were trying to design a disordered sequence to serve as a synthetic IDR linker between protein domains, one could upload a FASTA file of proposed disordered proteins with different lengths and compositions, predict the global dimensions of each, and then filter for a specific set of dimensions. This innovation greatly expands the potential for well-designed and controlled synthetic biology experiments and applications, and we are actively implementing such a design protocol into GOOSE.

In addition to the distribution through Google Colab, the ALBATROSS networks are also integrated within the SPARROW sequence analysis package under the “predictors” object operator. In this context, proteome-scale predictions can be achieved in a few lines of code on commodity hardware, e.g.:

```
from sparrow import Protein

# create a Protein object
P  = Protein(‘MEEPQSDPSVEPPLSQETFSDLWKLLPENNVLSPLPSQAMDDLMLSPDDIEQWFTEDPGPDEAPRMPEAAPPVAPAPAAPTPAAPA’)

# predict Rg
rg = P.predictor.radius_of_gyration()

print(rg)

> 32.381092
```

ALBATROSS predictions in SPARROW are, by default, memoized such that computations are not repeated after each call to the predictor operator. An optional override is provided to recompute predictions if desired. The lightweight and object-oriented nature of SPARROW makes it possible to build complex bioinformatic pipelines integrating both bioinformatic sequence properties as well as the emergent biophysical properties of a sequence.

In addition to performing prediction in series via SPARROW protein objects, users can also perform batch predictions on both CPUs or GPUs. If GPUs are available, predictions can be obtained at a rate of 1000s of sequence predictions per second. However, even on CPUs prediction provides offers 10-50x improvement in throughput with no loss of accuracy. As an example, $R_g$ values for all 29,998 IDRs defined by Tesei and Trolle *et al.* can be predicted in 23 seconds on a CPU using batch mode.

***Scaled vs. unscaled networks in ALBATROSS***

While scaled and unscaled networks were trained for end-to-end distance and radius of gyration (see above), scaled networks performed better across the board, but especially for short sequences. Notably, for sequences less than 30-35 residues in length, unscaled networks often predicted unphysical dimensions, while scaled networks performed uniformly well (**Fig. S17, S18**). Given the better performance of the scaled networks, the default networks implemented for sparrow predictions are the scaled networks. Moreover, for sequences shorter than 35 residues, even if unscaled networks are requested, sparrow falls back to force the scaled networks to be used. This behavior can be overridden by setting safe=False as a parameter when performing predictions.

***ALBATROSS comparison with expectations from the AFRC***

The Analytical Flory Random Coil (AFRC) – a Gaussian-chain-like model for disordered proteins – was compared to ALBATROSS to illustrate the breadth of chemistries where the gaussian-chain-like assumptions begin to break down[13]. We note that the derived radii of gyration from AFRC and predicted radii of gyration from ALBATROSS deviate in sequences with diverse sequence chemistries or patternings ($R^2$ = 0.676, **Fig. S19B**). The impact of sequence chemistry is more pronounced on the end-to-end distance than the radius of gyration, as illustrated by the weaker correlation between the ALBATROSS-predicted $R_e$ and the AFRC-derived $R_e$ ($R^2$ = 0.470, **Fig. S19D**).

***Notable errors in ALBATROSS***

Interestingly, we note that ALBATROSS performs less accurately on short synthetic sequences with large alanine sequence fractions (**Fig. S9C, D**). We note, however, that these alanine-rich sequences likely do not behave as *bona fide* disordered proteins. A similar trend was observed for the end-to-end distance network. Namely, shorter sequences with large alanine sequence fractions were more poorly predicted. Moreover, these challenges were largely mitigated via the scaled network training procedure. The utilization of the scaled networks for radii of gyration and end-to-end distance computations is thus the default setting, although we present an optional override for the use of the unscaled networks.

***Gene ontology enrichment***

Gene ontology (GO) enrichment was performed using PANTHER[16]. We calculated enrichment using all IDR-containing proteins as our background (using PANTHER Overrepresentation Test - Released 20221013). For all reported GO terms, we focussed on terms where (1) there were over 100 proteins with the term of interest.

***Comparison with CALVADOS2 radii of gyration***

Predictions for 29,998 IDRs calculated using the CALVADOS2 force field were obtained from Tesei & Trolle et al. [17], using the .csv file obtained from https://github.com/KULL-Centre/_2023_Tesei_IDRome/tree/main. We find that CALVADOS2 and ALBATROSS predictions correlated with an $R^2$ of 0.98 across the human proteome (Fig. S15) or 0.97 (for the $R_g$ values measured by SAXS). We also used the Google Colab notebook for CALVADOS2 simulations available at https://colab.research.google.com/github/KULL-Centre/EnsembleLab/blob/main/IDRLab.ipynb for our curated dataset of experimental radii of gyration from the literature.

### 2. Supplementary Figures

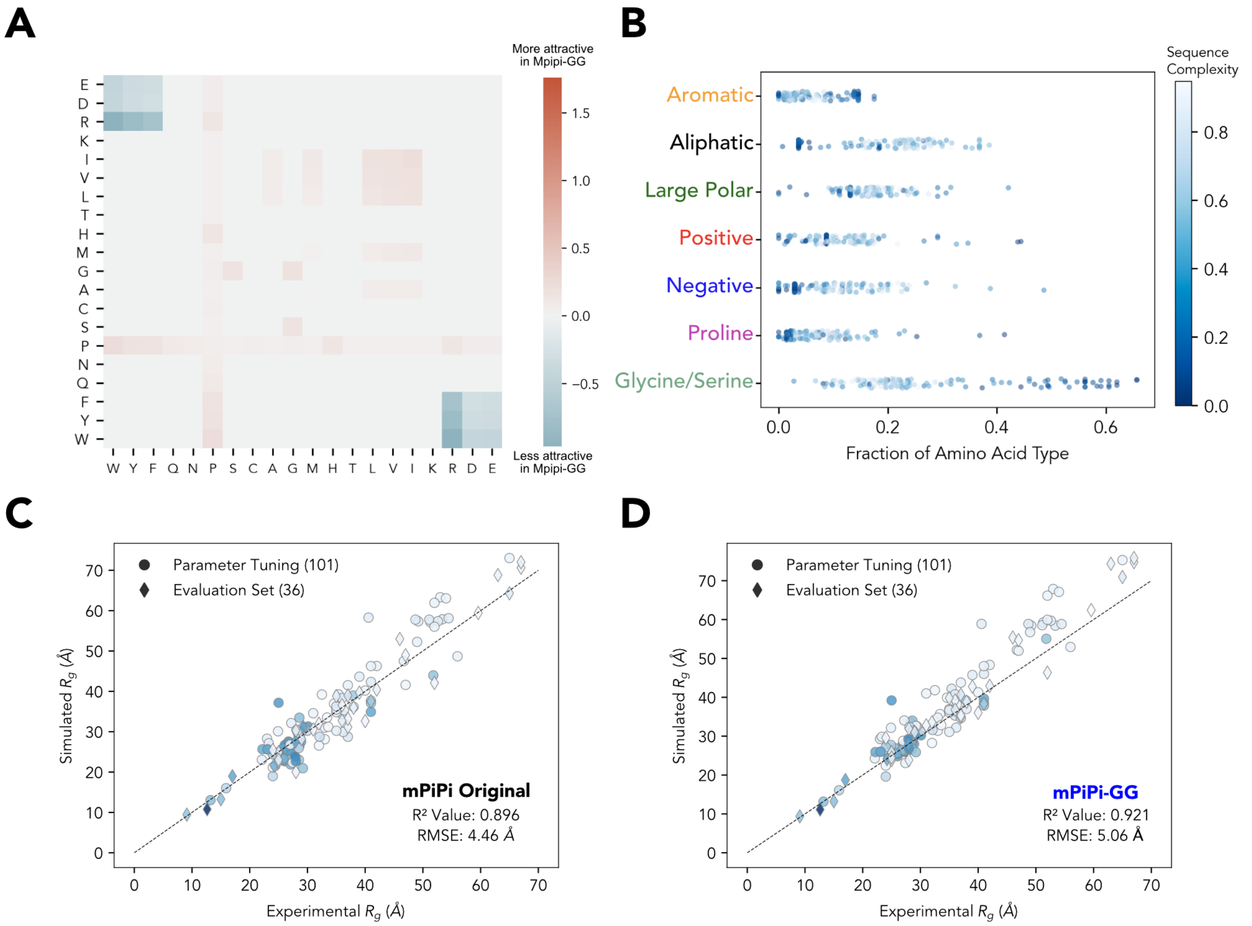


**Fig. S1. Reparameterization and accuracy of the Mpipi-GG force field. A)** Pairwise interaction matrix for the reparameterized Mpipi-GG force field. Pairwise interactions are colored by the relative change in interaction energies between the Mpipi force field and the new Mpipi-GG force field. Interaction energies are the sum of the net contributions from both the pairwise Coulombic interactions as well as the pairwise WF interactions. **B)** Composition of the curated experimental SAXS sequence dataset by amino acid type. The blue color gradient signifies the Wootton-Federhen complexity of the sequence. **C-D)** Correlations and RMSEs between the original Mpipi and Mpipi-GG force fields and a curated set of 137 experimental radii of gyration. 101 sequences (circles) were used for validating the Mpipi-GG force field, and 36 were new sequences held-out during parameter fitting (diamonds). Note that panel D is the same panel that appears in **Fig. 1C** (main text). The same color scale used in B is used in panels C and D.

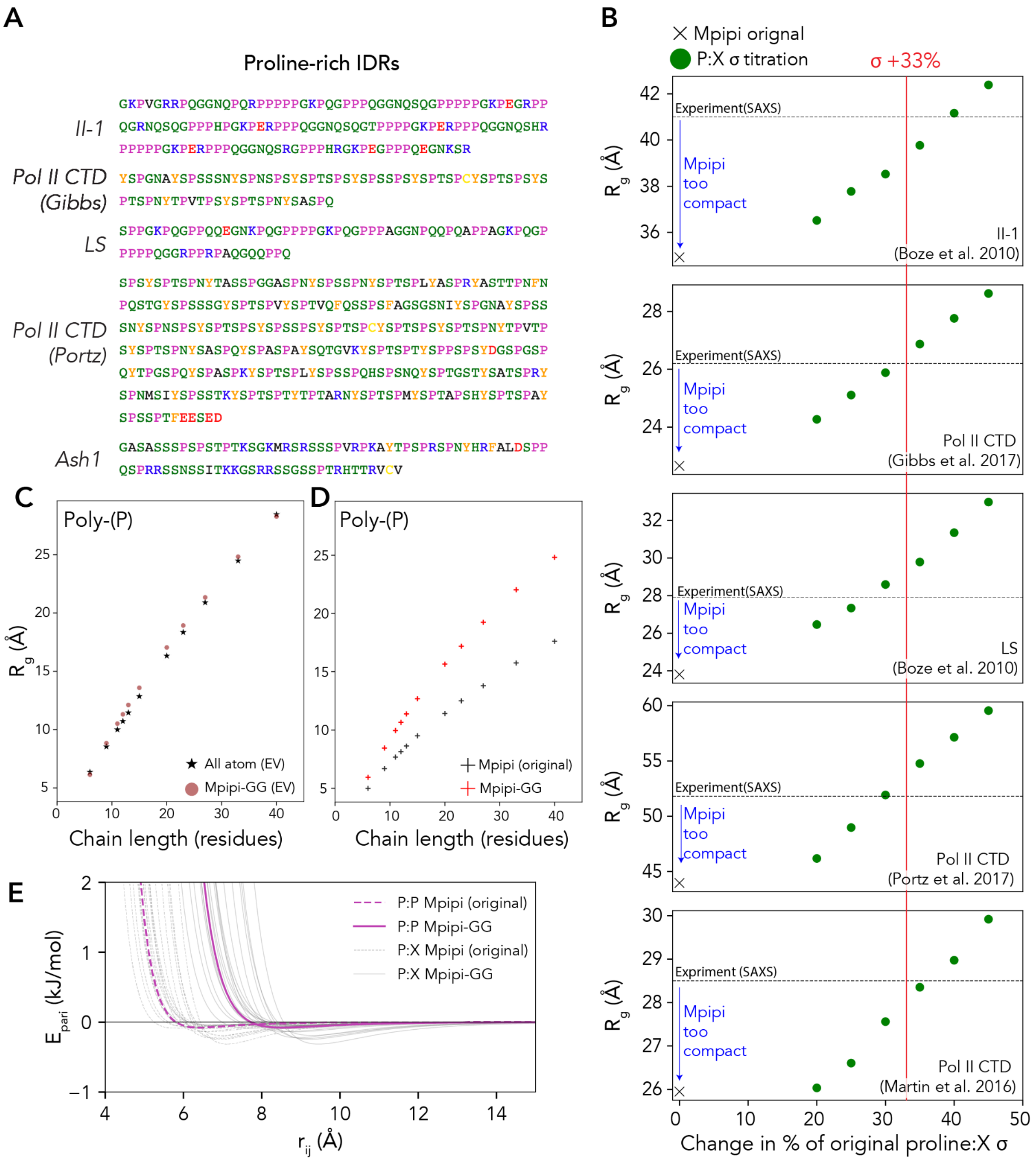


**Figure S2. Tuning of proline sigma (σ) parameter in the Mpipi force field**

**A)** Amino acid sequence of proline-rich IDRs previously studied by small angle X-ray scattering (SAXS)[3,5,18,19]. **B)** Comparison of experimental (dashed line) with predicted $R_g$ obtained from Mpipi original ('X' tick, far left) demonstrates that proline-rich IDRs are overly compact in Mpipi. This compaction can be alleviated by increasing the σ parameter from the WF potential (see equation 1), in effect, making proline residues larger. Green points represent the result of a systematic titration of the σ value. **C)** The optimal change to the proline σ parameters was selected by comparing excluded volume (EV) coarse-grained simulations from Mpipi with all-atom EV simulations and identifying the σ value that results in consistent $R_g$ vs. N scaling.

Shown here is a comparison of a +33% increase (used in Mpipi-GG) for Mpipi-GG EV simulations vs. all-atom EV simulations. **D)** Final comparison of polyproline dimensions for Mpipi-GG vs. original Mpipi. Mpipi-GG is more expanded than the original Mpipi, as also shown by the better agreement with experiment at a 33% increase, as shown in panel B. **E)** Wang-Frenkel (WF) potential for Pro:Pro interaction in the original Mpipi parameters (dashed purple line) vs. Mpipi-GG (solid purple line). Dashed and solid gray lines represent proline and each of the other twenty amino acids for Mpipi and Mpipi-GG, respectively.

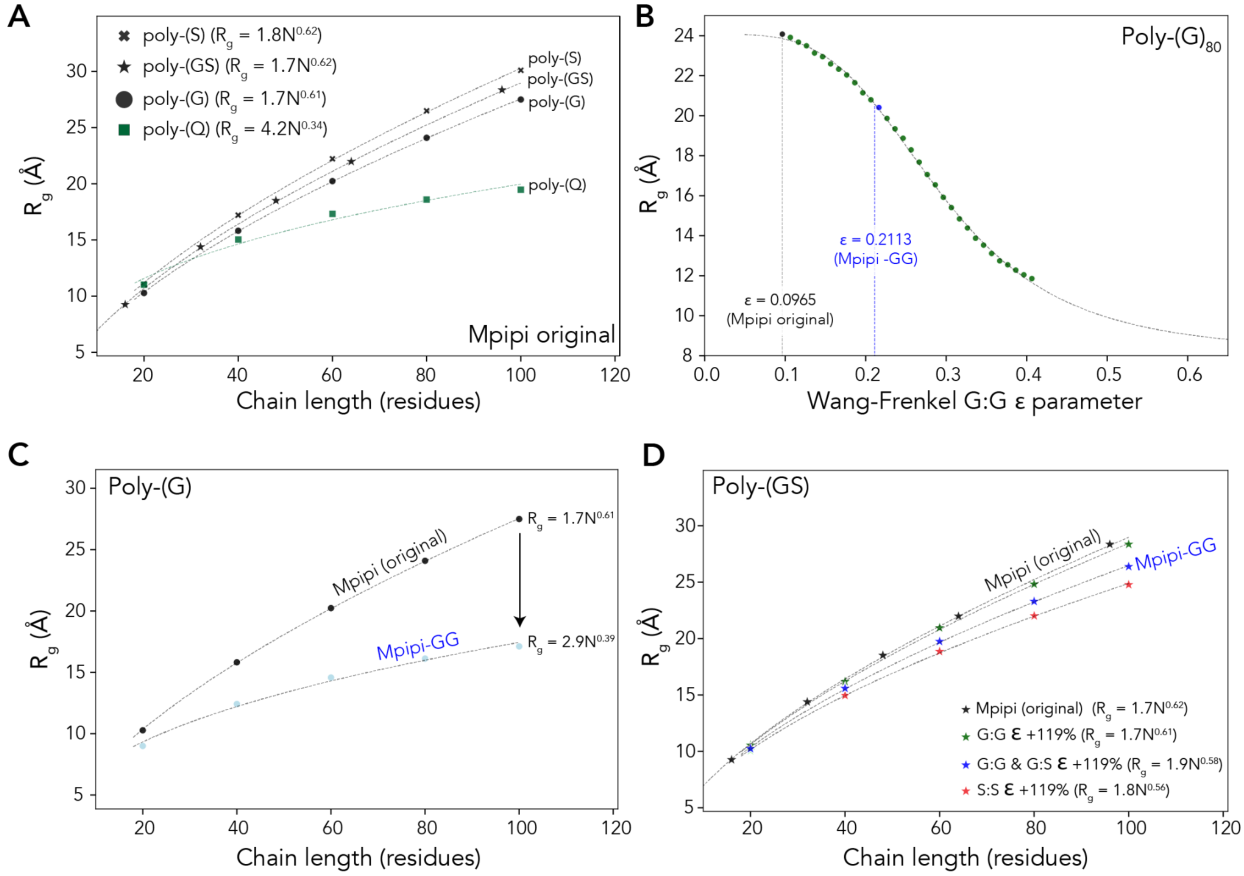


**Figure S3. Gly/Ser Mpipi-GG reparameterization A)** Simulated scaling behavior for simple polymeric sequences performed using the original Mpipi model. Poly-(Q) compaction is consistent with experimental work[7]. However, poly-(G) scales as a self-avoiding random chain, despite prior work implicating a scaling exponent closer to 0.40[11]. Further, poly-(GS) scales as a self-avoiding random walk ($\nu$ = 0.6) against prior work from simulations and experiments, which suggest poly-(GS) sequences behave closer to a Gaussian chain ($\nu$ = 0.5 - 0.55) [7–10]. These data suggest that G:G interactions are too weak. **B)** To reparameterize G:G strength we systematically titrated the glycine ε parameter, leading to a coil-to-globule titration from which we selected the ε value that best matches the expected scaling of 0.4. **C)** Comparison of Mpipi vs. Mpipi-GG, revealing the more compact scaling and smaller scaling exponent (0.39 vs. 0.61), in better agreement with experiment. **D)** Despite strengthening G:G interactions, poly-(GS) dipeptide repeat polymers are still relatively expanded (compare black and green data). To address this, we asked how changing the S:S: interaction (red) vs. G:G and G:S (blue) altered chain dimensions. Given the prevalence of serine residues in disordered regions, we released the Mpipi S:S interaction strength was likely already reasonable, such that we selected the same scaling for G:G and G:S to enhance cohesive interactions between glycine and serine.

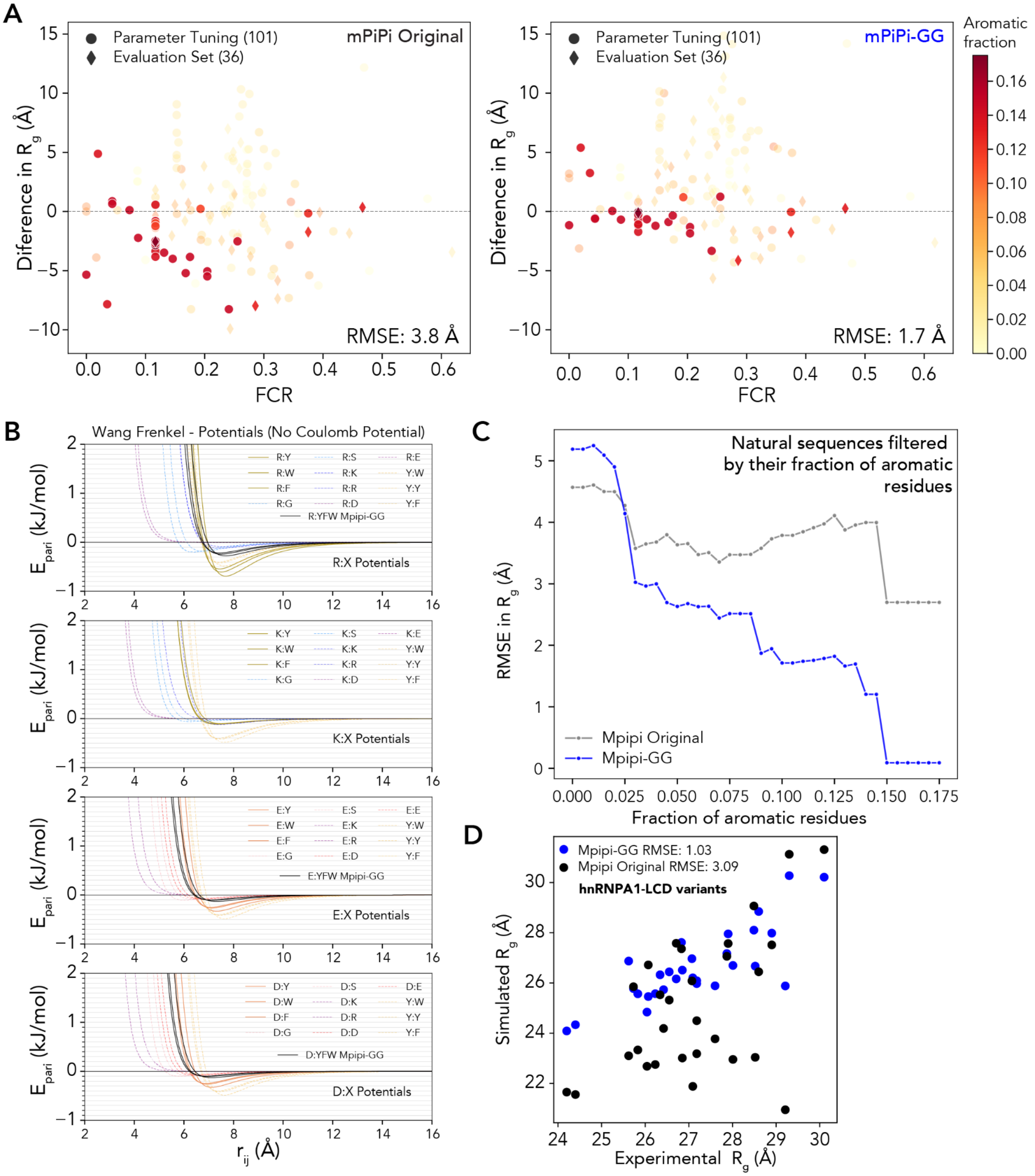


**Figure S4. Reparameterization of pairwise aromatic interactions for the Mpipi-GG force field. A)** Residual plot for the deviations between experiment and Mpipi (left) and Mpipi-GG (right) as a function of the fraction of charged residues. When comparing all sequences in our curated dataset with >10% aromatic residues, the original Mpipi has an RMSE of 3.8 Å, whereas Mpipi-GG has an RMSE of 1.7 Å. We note specific improvement in sequences that are jointly aromatic and charge rich - i.e., >10% of charged residues by fraction. **B)** Pairwise Wang-Frenkel interaction potentials for arginine, lysine, aspartic acid, and glutamic acid relative to aromatic residues (solid lines) and benchmark residues (dotted lines). Updated Mpipi-GG potentials are drawn in black. **C)** Panel **A** uses an aromatic threshold value of 0.10; however, this choice is somewhat arbitrary. Therefore, we chose to demonstrate generality by looking at many different potential thresholds. This plot looks at the root mean squared error as a function of aromatic thresholding. For each threshold value (x-axis), we took all sequences in the curated dataset with aromatic amino acid fractions equal to or exceeding the respective threshold value and computed the RMSE between the experiment and simulated results for each respective force field. As aromatics fractions are increased, Mpipi-GG consistently has modest improvements in recapitulating experimental SAXS radii of gyration. RMSEs near zero in Mpipi-GG are reflective of the fact that there are few sequences with greater than 15% aromatics in the curated library. Nevertheless, Mpipi-GG is highly accurate for these aromatic and charge-rich sequences. **D)** Comparison of simulated $R_g$ vs. SAXS-derived $R_g$ for hnRNPA1-LCD variants[20,21]. These sequences systematically vary charge and aromatic content, providing a convenient reference set for comparing Mpipi-GG vs. Mpipi in the context of aromatic/charge interactions.

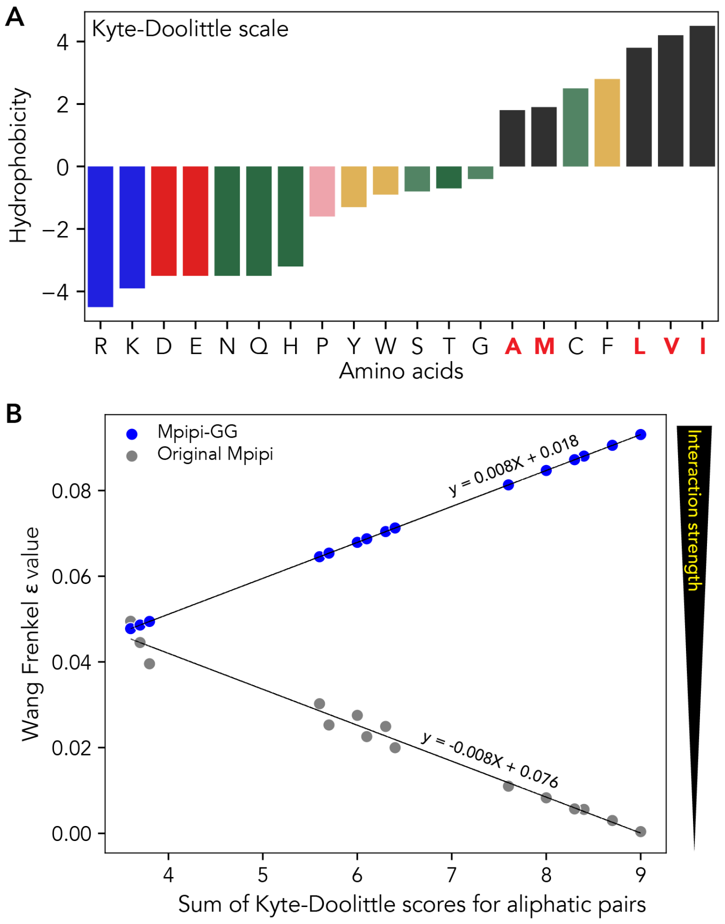


**Figure S5. Reparameterization of pairwise aromatic interactions for the Mpipi-GG force field. A)** Kyte-Doolittle hydropathy scale for each of the twenty amino acids. **B)** Pairwise sum of Kyte Doolittle hydrophobicity ($KD_{hyro}$) values relative to Mpipi $\varepsilon_{AMLVI,AMLVI}$ values. Reparameterized $\varepsilon_{AMLVI,AMLVI}$ values in Mpipi-GG, $\varepsilon_{i,j}$ are equal to 0.0008*$KD_{hyro(i+j)}$ +0.018, where the slope of this line is inversely proportional to that of the $\varepsilon_{AMLVI,AMLVI}$ values in the original Mpipi force field, but scaled so the more hydrophobic pairs are stronger, as opposed to weaker.

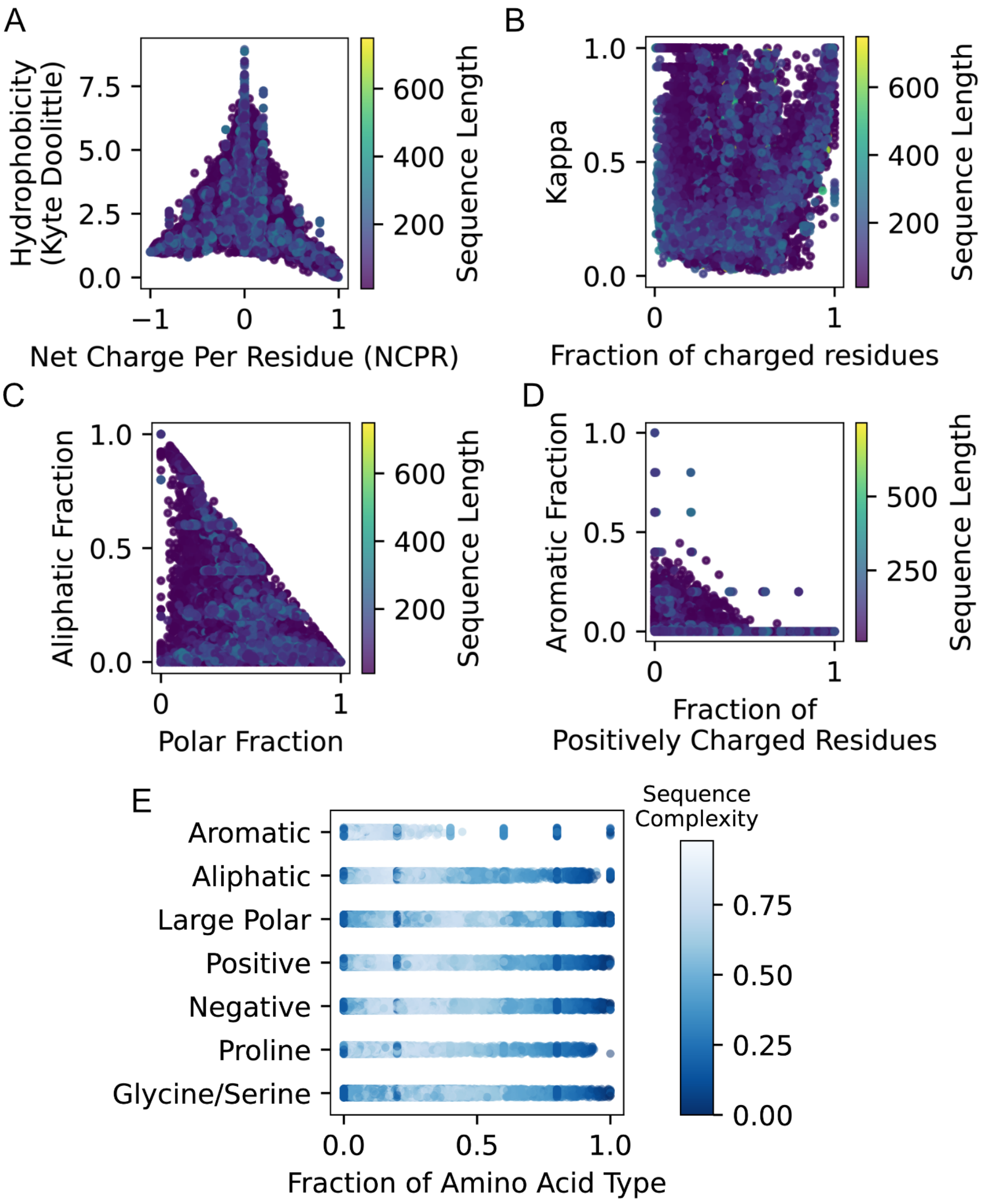


**Fig. S6. Composition of the IDR sequence library used for training. A)** Composition of the synthetic sequence library by amino acid type. The blue color gradient signifies the Wootton-Federhen complexity of the sequence. Two-dimensional scatter plots showing the chemical space explored by our synthetic IDR library. Each point in all panels is colored by the length of that particular sequence. **B)** Fraction of polar residues versus the fraction of aliphatic residues in a given sequence. **C)** Fraction of aromatic residues and the fraction of positively charged residues (RK). **D)** Fraction of charged residues versus the charge patterning parameter kappa ($\kappa$).

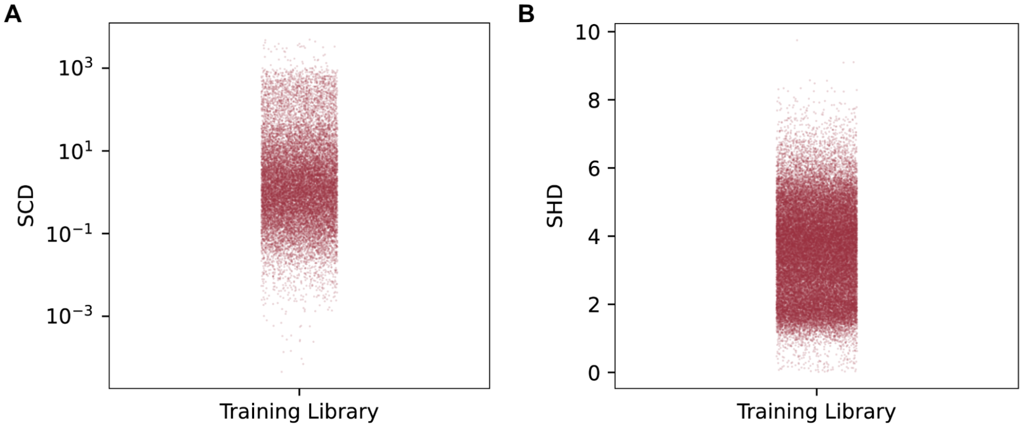


**Figure S7. Sequence property comparison for the biological and synthetic sequence libraries. A)** Distribution of sequence charge decoration (SCD) values for the training set of sequences. Note the SCD is plotted on a logarithmic scale as the training data covers a broad dynamic range of SCD values. **B)** Distribution of sequence hydropathy decoration values for the training set of sequences.

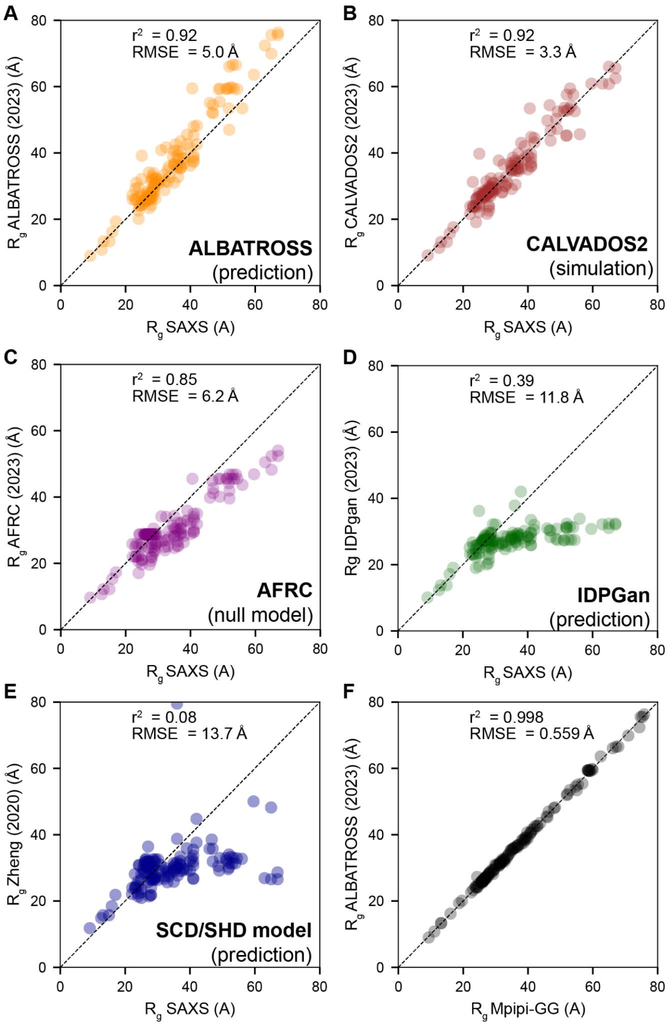


**Figure S8. Comparing predictions vs. SAXS measurements for state-of-the-art tools for mapping between sequence and ensemble. A)** The ALBATROSS $R_g$ network recovers a correlation coefficient of 0.92 (RMSE = 5.0 Å), in agreement with analogous analysis for Mpipi-GG. Predicting these values in series takes ~4 seconds on a CPU. **B)** Simulations performed using CALVADOS2 show an equivalent correlation coefficient with a smaller RMSE value (RMSE = 5.0 Å)[17]. Simulations performed using the Google colab notebook took ~ 7 minutes on GPU for a 137 residue sequence. **C)** The AFRC is an analytical model that reports the expected dimensions of a polypeptide if it behaves as a Gaussian chain[13]. Calculating anticipated $R_g$ values for these sequences takes < 1 second for all sequences. **D)** IDPGan is a deep learning model trained to predict IDP ensembles from sequence[22]. Predictions were performed using the Colab notebook and took 2-5 seconds per sequence. **E)** The SCH/SHD model uses a set of empirical equations derived by Zheng *et al.* to predict the radius of gyration from sequence via a predicted scaling exponent[23]. Predictions take < 1 second for all sequences. **F)** ALBATROSS reproduces simulation-derived radii of gyration with almost no appreciable error.

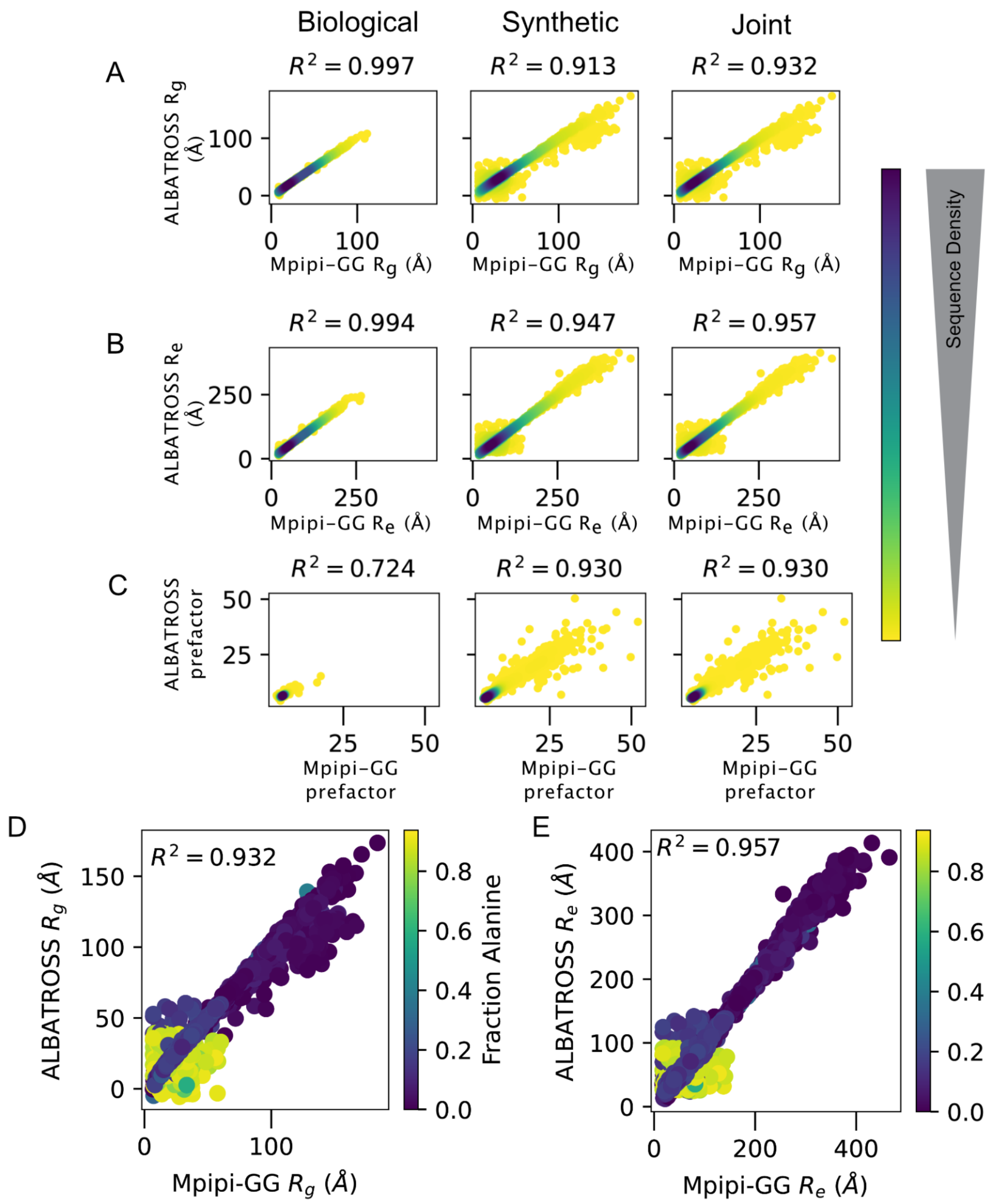


**Figure S9. Evaluating the performance of the ALBATROSS networks on held-out test sets A-C)** Performance of the unscaled radius of gyration network, unscaled end-to-end distance network, and polymeric prefactor for biological, synthetic, and the joint test set. **D-E)** Examining the unscaled radius of gyration and end-to-end distance ALBATROSS networks. We note that for these networks, shorter sequences with large alanine sequence fractions display predictions with large variations from the true simulated Mpipi-GG values. For this reason, we default to using the scaled networks trained as presented in **Fig. 2.** Data points are colored by the alanine sequence fraction.

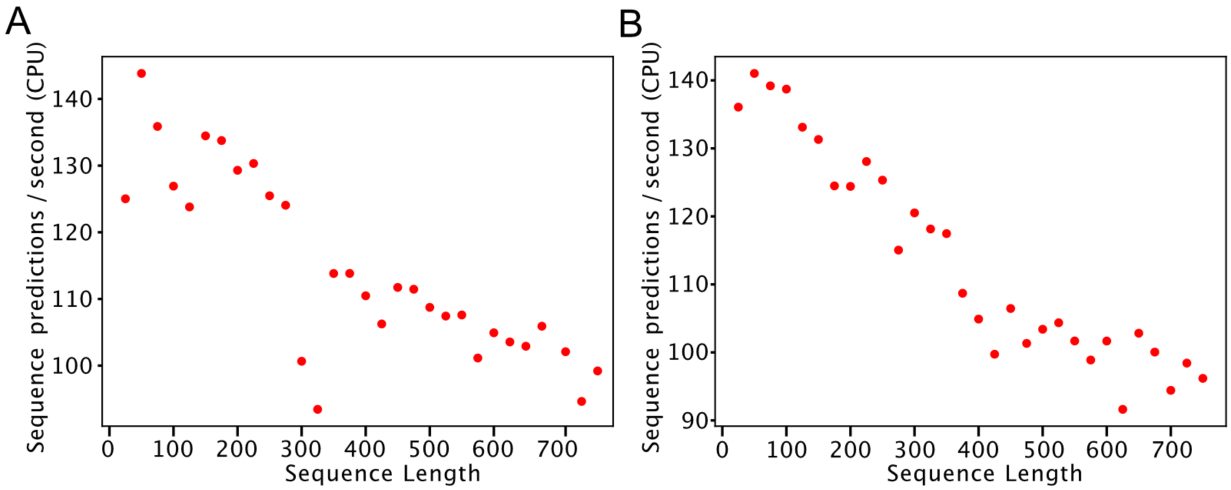


**Figure S10. Network performance on standard commodity hardware.** We measured predictive power on standard CPUs for **A)** the radii of gyration network and **B)** the end-to-end distance network as a function of sequence length. For 100-residue IDRs, performance sits around 120-140 sequences per second. We emphasize that on a Google Colab notebook using GPUs, the entire human proteome takes ~8 seconds, but we focus here on CPU performance given the broad availability of CPUs.

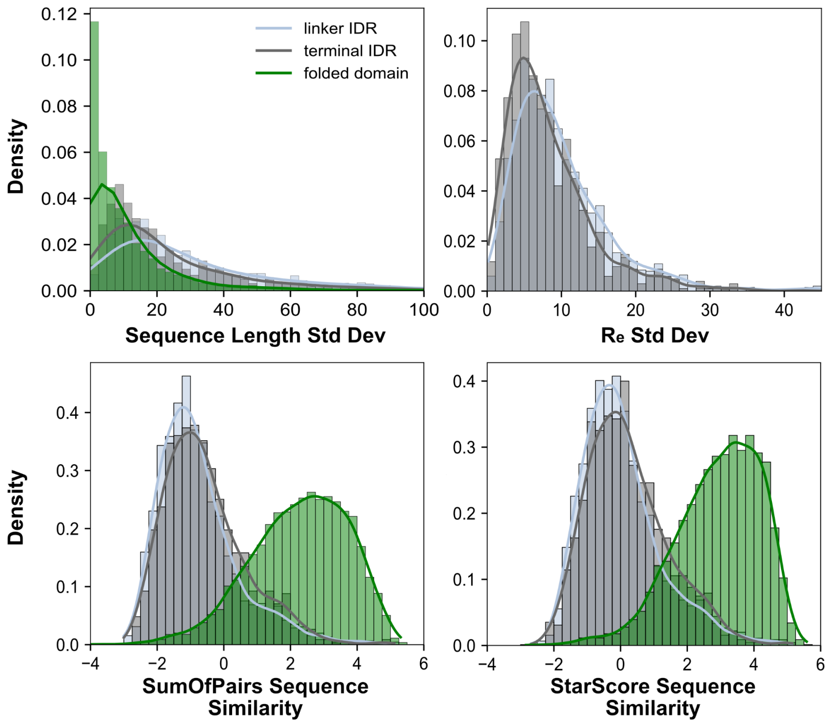


**Figure S11. Distribution of predicted $R_e$ standard deviation and sequence similarity standard deviations across yeast homologs.** Histograms describe the scatterplot data presented in **Fig. 6B** and **S12**. For sequence similarity metrics, the distribution of homologous folded domains is plotted for comparison.

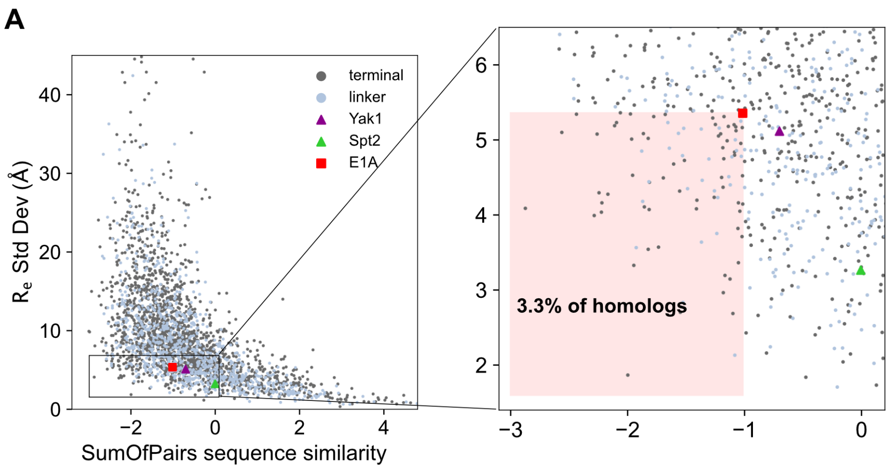


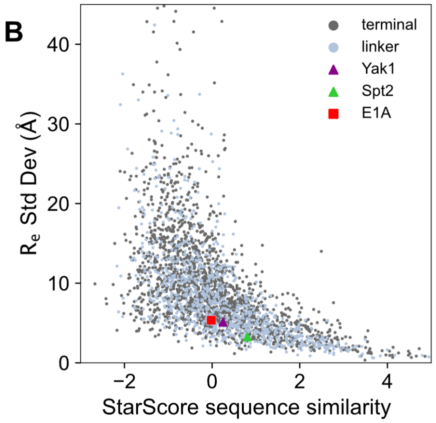


**Figure S12. Multiple sequence alignment-based sequence similarity analysis of yeast IDR homologs. A)** Related to **Fig. 6B**; sequence similarity is calculated via the SumOfPairs method (pyMSA package) on the IDR multiple sequence alignment. Negative values correspond to more divergent primary sequences. There is a trend that more divergent homologs tend to have a greater variation in predicted $R_e$. The panel on the right shows a zoomed-in inset of the region of the plot corresponding to homologs that have more divergent sequences, and more constrained $R_e$ than E1A. **B)** Same plot as **A**, but with sequence similarity calculated using the StarScore method, indicating this analysis is robust to the choice of specific metric.

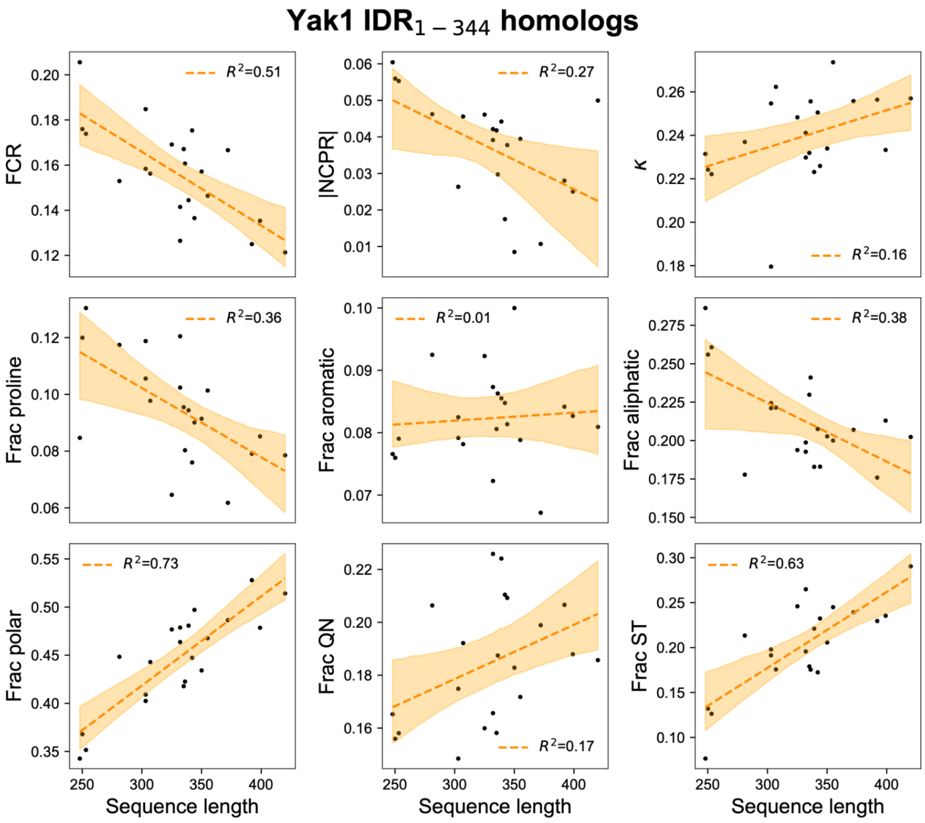


**Figure S13. Expanded comparison of Yak1 IDR homologs' sequence features.** Yak1 homolog sequence length compared to various protein sequence features. Line of best fit and 95% CI denoted in orange. FCR=fraction of charged residues; |NCPR|=absolute value of net charge per residue.

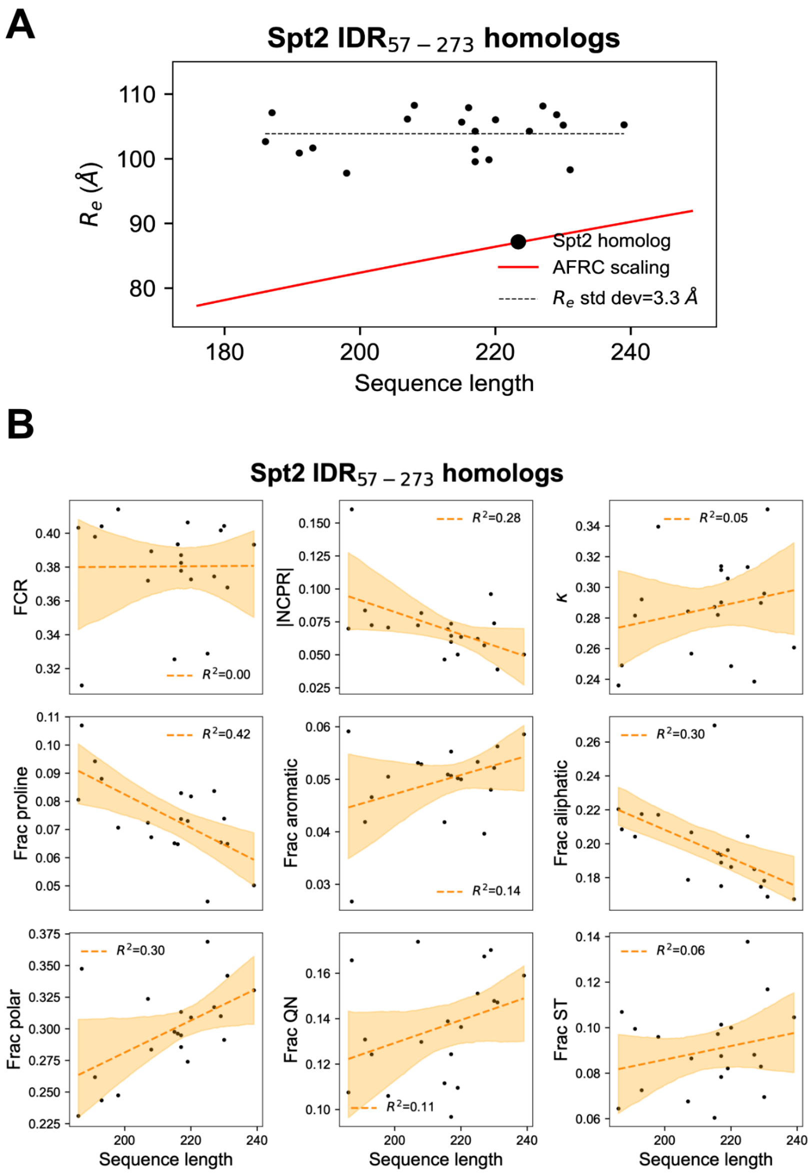


**Figure S14. Spt2 IDR has conserved dimensions across homologs despite divergent sequences. A)** Sequence length and ALBATROSS-predicted $R_e$ for Spt2 and its yeast homologs. The gray dashed line denotes the mean $R_e$. The red line denotes $R_e$ as a function of sequence length for an Analytical Flory Random Coil polymer scaling model. **B)** Spt2 homolog sequence length compared to various protein sequence features. Line of best fit and 95% CI denoted in orange. FCR=fraction of charged residues; |NCPR|=absolute value of net charge per residue.

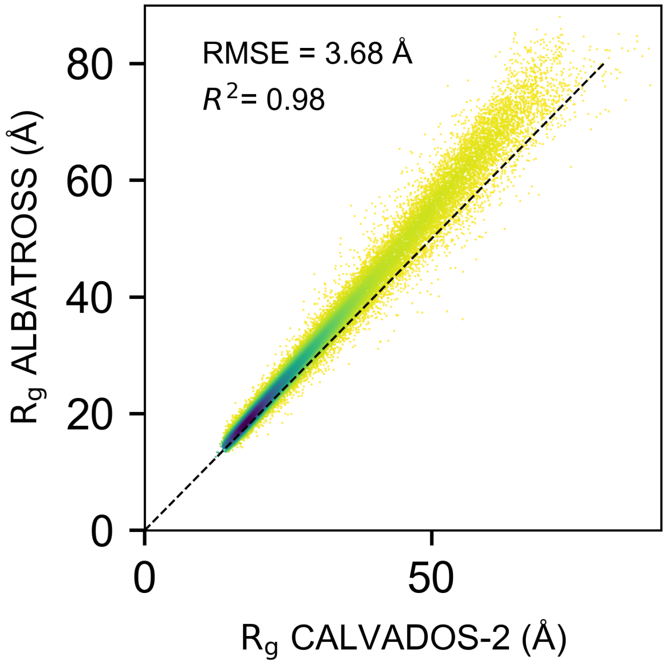


**Figure S15**. **Comparison of ALBATROSS predictions and $R_g$ values obtained from Tesei & Trolle *et al.* IDR-ome**[17]. The root mean square error here is equivalent to the mean square error between Mpipi-GG and SAXS data (see **Fig. S8**). ALBATROSS $R_g$ values are systematically slightly more expanded than CALVADOS2 values, a difference that is in part expanded by the proline-drive expansion seen in Mpipi-GG and hence ALBATROSS. However, this may also reflect a slight underestimation of hydrophobic interactions in Mpipi-GG and hence ALBATROSS.

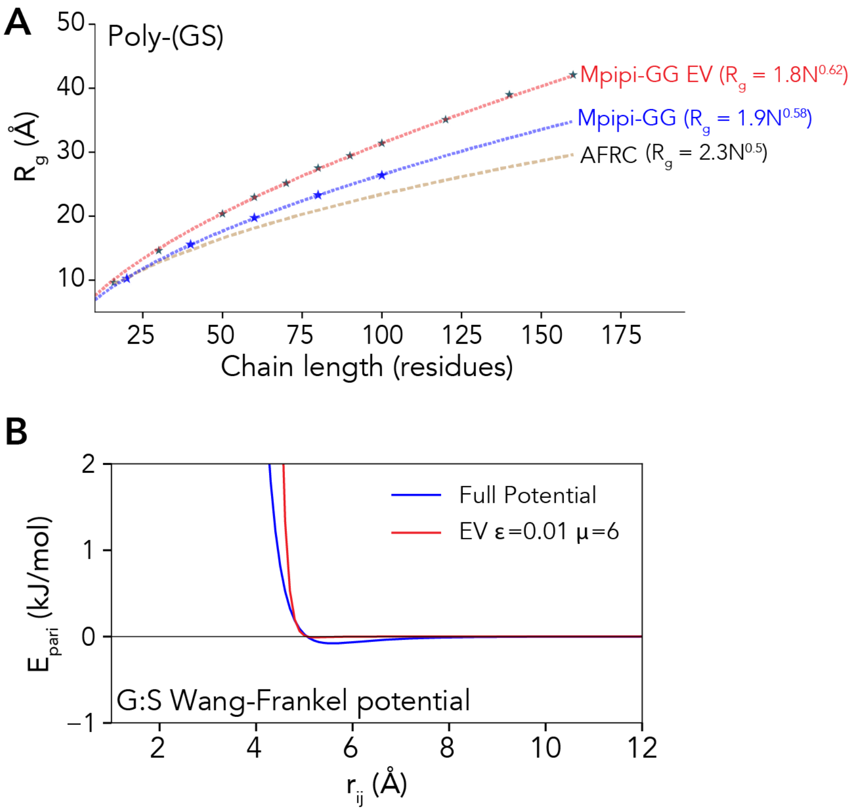


**Figure S16. Parameterizing an excluded volume model for the Mpipi-GG force field. A)** Scaling behavior for poly-GS as represented in terms of Mpipi-GG (blue) Mpipi-GG as an excluded volume (EV) simulation, and the Analytical Flory Random Coil (AFRC) model. EV simulations are more extended than full Mipi-GG simulations. **B)** Wang-Frankel interaction potentials for a representative pair of beads (G:S). The full Wang-Frankel potentials for the interaction of glycine and serine in the Mpipi-GG forcefield - note the dip near $r_{ij}$ of ~5.5, reflecting the attractive part of the potential. After tuning the $\boldsymbol{\sigma}$ and $\boldsymbol{\mu}$ parameters, we obtained a pairwise interaction potential with near zero attractive interactions (red) that match the same dimensions as the full Mpipi-GG forcefield.

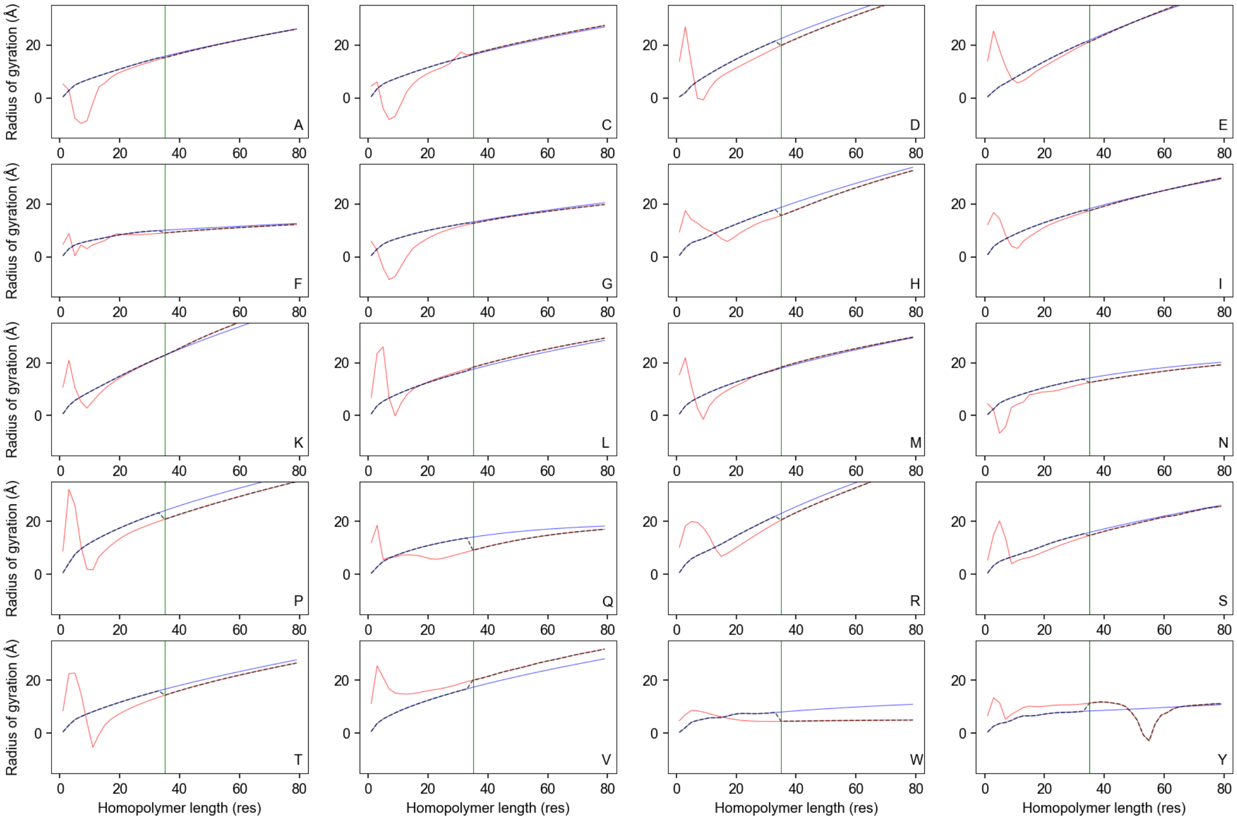


**Fig. S17**. Comparison of the scaled radius of gyration networks (blue) vs. non-scaled radius of gyration networks (red) for a set of 20 homo-polymeric sequences. Above a length of 35 amino acids (green vertical line), reasonable agreement between the two models is obtained, with some exceptions (notably poly-tryptophan and poly-tyrosine). However, below this threshold, we often see substantial deviations between the scaled networks and non-scaled networks, with non-scaled networks showing unphysical behavior. Based on these observations, in ALBATROSS V2 we use a threshold of 35 residues and, by default, even if a non-scaled network prediction is requested, we fall back to using the scaled network prediction to avoid nonsensical $R_g$ and $R_e$ values. Encouragingly, the agreement between the two models above 35 amino acids is much closer for non-homopolymeric sequences (see **Fig.S18**).

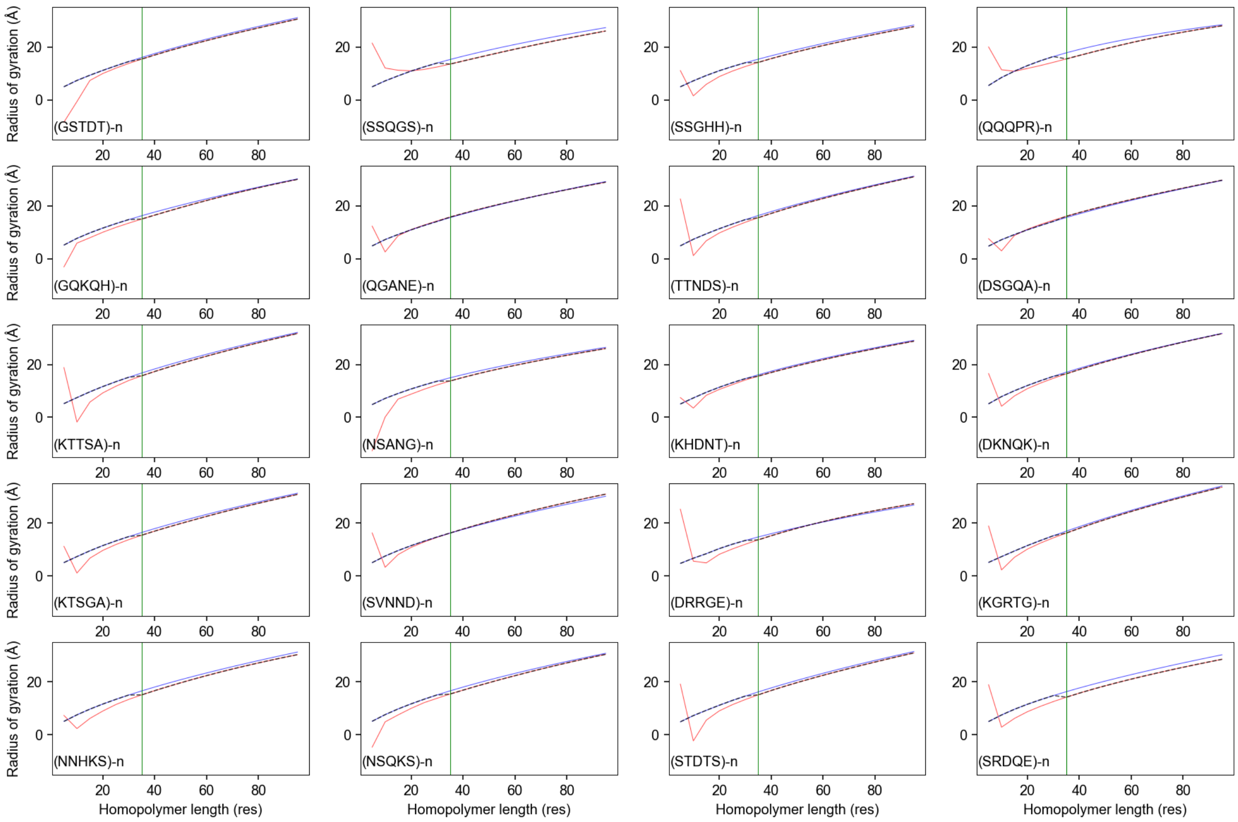


**Fig. S18**. Comparison of the scaled radius of gyration networks (blue) vs. non-scaled radius of gyration networks (red) for a set of 20 random disordered repeat proteins. Above a length of 35 amino acids (green vertical line), good agreement between the two models is obtained, but below this line, we often see substantial deviations, where the non-scaled network (red) shows unphysical behavior.

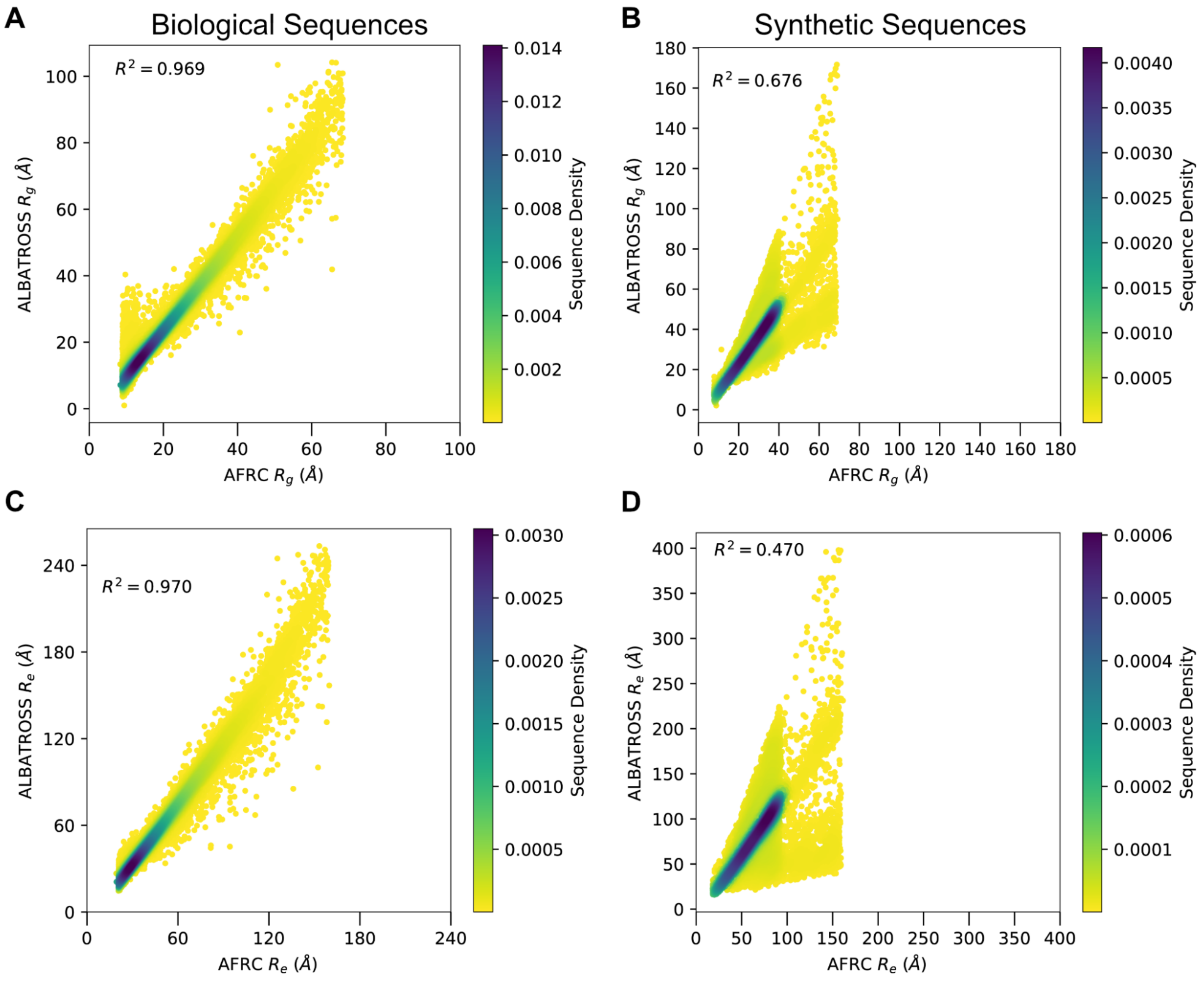


**Figure S19. Comparing the Analytical Flory Random Coil (AFRC) chain dimensions and the ALBATROSS predicted chain dimensions for the synthetic and biological sequence libraries. A-B)** Correlations between modeled radii of gyration for both biological sequences (right) and the synthetic sequences (left). **C-D)** Correlations between modeled end-to-end distances for both biological sequences (right) and the synthetic sequences (left).

### 3. Supplementary Tables

**See separate file:** *final_GO_analysis_compact_IDRs.xlsx*

**Table S1. Gene ontology analysis for proteins with compact IDRs.** For annotations that contained 100 or more entries in the basis set and showed 2-fold or higher enrichment, the vast majority of annotated term pertains to RNA in some way across the three classes of gene ontology.

**See separate file:** *final_GO_analysis_expanded_IDRs.xlsx*

**Table S2. Gene ontology analysis for proteins with expanded IDRs.** For annotations that contained 100 or more entries in the basis set and showed 2-fold or higher enrichment, a variety of IDR-associated annotations are identified, including chromatin binding, cytoskeletal regulation, and cellular organization, in good agreement with analogous analysis from Tesei & Trolle, despite the two analyses being done in different ways [17].

| Organism | ID | Res. # | Sequence | Pred. $R_e$ (Å) |
|---|---|---|---|---|
| *S. cerevisiae* | YJL141C | 1-344 | MNSSNNNDSSSSNSNMNNSLSPTLVTHSDASMGSGRASPDNSHMGRGIW<br>N<br>PSYVNQGSQRSPQQQHQNHHQQQQQQQQQQQQNSQFCFVNPWNEEKVTN<br>S<br>QQNLVYPPQYDDLNSNESLDAYRRRKSSLVVPPARAPAPNPFQYDSYPA<br>Y<br>TSSNTSLAGNSSGQYPSGYQQQQQQVYQQGAIHPSQFGSRFVPSLYDRQ<br>D<br>FQRRQSLAATNYSSNFSSLNSNTNQGTNSIPVMSPYRRLSAYPPSTSPP<br>L<br>QPPFKQLRRDEVQGQKLSIPQMQLCNSKNDLQPVLNATPKFRRASLNSK<br>T<br>ISPLVSVTKSLITTYSLCSPEFTYQTSKNPKRVLTKPSEGKCNN | 111.9 |
| *V. polyspora* | 1060.53 | 1-303 | MQGEVMSNNRPFDNGNHNGNSNNNEEKKDFWNPAFMTQQRGFQQQQQQQ<br>P<br>QQSALNSDIDPGMRPGQFIFNNPWRNDPASRKNSLAASPFPDNSMGMLD<br>I<br>DYRRRKSSLVTPPSRTANTNPFQSDKAANFPQSTAAYQQQRQFLANDNG<br>M<br>FADRLGPSMNFVDPNDFYRRQSVAAVSYQPDTGPTITANQSIPITHYRR<br>L<br>SAYPQSMTPSLHPIRREEPLVKQQPIHIPQMHITNSANDLQPLINPTPD<br>Y<br>RRASLNSKEISPLNALTTGLITTYSLCSPDFAYKTSKNPKRVLTKPNEP<br>R<br>GNN | 111.3 |
| *V. polyspora* | 2001.10 | 1-350 | MKADSSSNTNDPNSDNENDKQNMWNPIFINQNPVRYMSQQQQQHQYQHL<br>P<br>QNLQHQQQQQLTQEYSNSQQMAFVNPWMAATDDSINYSPLYSTAPPSTL<br>S<br>VLQESDEHPQKIPCFDNYNYDQRRKSSIIIPPTREPAPNPYLQEYYNGY<br>P<br>APVFQLETNEMMNQQQQYNPNYFICNQPLYQQSGDSTFSIADPTMSFSN<br>Q | 112.3 |

| | | | | |
|---|---|---|---|---|
| | | | DLTMRQSVGPEHFISDHTNSNQYSKNGKSTQQLISYRRLSAFPQTNGFH<br>T<br>LQPPMYISSEYRKSSMSVSPKPLMISHLKQCHSKADLEPVMNQTPKFRR<br>A<br>SLNSKTISPLIALTKGLITTYSLCSPSFSYKTCKNPKRVLTKPSDGKFN<br>N | |
| *T. phaffii* | TPHA0A0457<br>0 | 1-325 | MEVDNEEISNGGGFWNPKYLEQNDSNGSGGNNNNNNDNNNSQGIQWNQN<br>F<br>LDSQPNVAASQVNSNFNMKRRKSSIIITPPSRTANTNPFQNKNIISDGN<br>Y<br>NNSYIRNYDTTRRGSIAGYNYSHVSNNSVRFPQQQLAPQQQQQSYYNQP<br>P<br>LGGYSLQKNLHRSKVKGFIDPNDLQRRQSVATVNYDTKQHINNINPNTG<br>K<br>PSVQNLHGSFNFRRRQSAYPIFNDSFAYNNSSGGNKKFSIQKNDPINLI<br>P<br>TMFNVQSKSDLKPTLNILKDDTRRASINSQKISPLHALTSGLVTTYSLC<br>S<br>PDFQYNLSKNPKRVLTKPSEPAYNN | 103.5 |
| *T. blattae* | TBLA0B06510 | 1-342 | MENQTIQEYPENNSEEGAPNDTSYKTPLQHRGSQSYSHQQLHNGQRNSL<br>T<br>FINPWGSNLNGSSDLFMQSNQSLLSTSSENQRRPSMDDFKRRKSSIVIP<br>P<br>SRAPGINPYFYNIDTTATFNGDGTSDNFQQNQNIDANQQQKKIFADEKD<br>L<br>YVQALLDPSSGIYDPDAFYRRRSLAAPAFTSSMGSSSMNTANNTATTNA<br>S<br>SYQLNEPISPSNVNFSLMPHGCGINSRQPNRTGSFRKLSAYGAPVQSTF<br>K<br>STYREEYNLLQQPPLEIPQMTPCNSKSDLQPTLNKLPKYRRASLHSSTI<br>S<br>PLVGLTKGLITTYSLCSSDFQYKTSKNPRRVLTKPNEGVANN | 120.9 |
| *T. blattae* | TBLA0D05770 | 1-372 | MQRRYSELDPEDMDINSRSNSNSSSNSLHNNAMTTLMQESSTKMQFKNQ<br>H<br>RFNNPWGNGSNNNNDTFDYLGSRKEFDSMEHHVSIMEPEELTGNNSVGS<br>I<br>TCSDINNEQYKRRKSSVIIPPTREAGPNLYQHLMNKNWQATSNNNGNNN<br>N<br>LNMNNNLFQNMQPNSTTTSNVNISLNMNTNMNFHMQNPDVNSRFLANED<br>S<br>FRRQSLATFNYNPQHTATNLHSHYNPNMVNSANPTNNNVLLRNPSTNST<br>T<br>TSPYRRMSTFPQNINPLASFNNQVLKDENQLHLLQQSQSQSQQPIIIPH<br>M<br>TKCTNKSDLQPIINDQPEYRRASTFTNNVSPLKSLTTRLITTYSLCSKD<br>F<br>IYKTSRNPKRVLTKPCEPVEND | 119.7 |
| *N. dairenensis* | NDAI0G06240 | 1-420 | MESIHNGTMSSNGNAAVQITNNDNNLKNNSPFNNSIWKANYVENSQQQQ<br>Q<br>QQQQQLQSKDSLTPGGVSPARKQQGMGPPSSIPQQQQQQQQQQFSFTNP<br>W<br>NKNDTVLPQQTYNYAMDEDMLESFKRRKSSLVIPPSRARAPNPFEFNFQ<br>Y<br>QAFPQQSSQQGINNNSNSNPYSYQQYQQHQLYNTHQYNNPNLMAANGNN<br>N<br>SRFFNQQDFHRRQSVAASQFYPSTNTSNISNNTKSMMNSPNSVSMQSNN<br>N<br>NIHSSNTYLPTATPTNTSNNNIMVNNNNNNNNNNNNNNNNNYNPNSAIT<br>A<br>SFRRLSAYPTTGISSTIPSQQQLLRKGSILEASPSAAATAQPIVIPTMK<br>K<br>VNFKKDLKPIINPTPKFRRASLNSKTISPLLALTRNLITTYQLCSPDFT<br>Y<br>KPSKNPKRVLTKPSEGKFNN | 121.2 |

| | | | | |
|---|---|---|---|---|
| *N. castellii* | NCAS0I02680 | 1-355 | MTTPNDFASPKEMSPRHANSSESLHNNHQLQQGAADQANNNNNIWKSTF<br>M<br>NQQQQQQQQQNSFAFTNPWSSNNADSTLSPPQQQQQSYNMDMDEDMLES<br>F<br>KRRKSSLVIPPSRAPAPNPFQYDQYNQYGPPQSQQQVPSQQQEINYSHQ<br>Q<br>QQQLYNAHQYGARLGPSMYQDFQRRQSMAASHLYPNGNGNNITNYSTRT<br>I<br>NKNSSITPNTANPSMMAGPSYIPGGLSNDNSMVSPFRRLSAYPTTNIHT<br>P<br>ATANLQPPYKQLRRTGEEQQQQQQPKPLIIPSMKVCKNRNDLNPITNPT<br>P<br>KFRRASLNSKTISPLLALTKNLITTYQLCSPDFTYKPSKNPKRVLTKPS<br>E<br>GKFNN | 118.2 |
| *K. naganishii* | KNAG0C0260<br>0 | 1-392 | MHQNSSQMTNNNGNSIWNPNFFQNDEGQQQQQQEQQQQQQQQQKQEVKF<br>A<br>FNNPWDSNGGNQNDYSTGQVAGDAAGNTGHVLYSPNHEMIMETPNEDFE<br>N<br>FKRRKSSLVIPPTRAGAPNPFQYENYPTYNSMGQNTSSINSNGDYGPGG<br>A<br>SSSAGPGTALSGFNPSMGSSQQQRQSLFNKQQQQQQHLNTRFVPNLYNQ<br>Q<br>NGFQRRQSMAATAFTTPQYTTAPSTAGNTTSNFGNNAGTVGPNGHFSTN<br>M<br>ANSSSAGKLQNVQSGMNPNFMGMSSVSPYRRLSAFPTSSGGMTSATPTT<br>S<br>TPFPQQFAPLREEKPTIPQMSSCKSKSELSPTVNKTPKYRRASVHSQTI<br>S<br>PLNAMTKNLITTYQLCSPDFIYKTSKNPKRVLTKPSEGKCNN | 119.1 |
| *K. africana* | KAFR0F0194<br>0 | 1-399 | MDQTNGQQMNSPFDNSGNTNNSTNNTNNNTNNNGNNDGSDNGYHLWNQD<br>Y<br>LSPGHPQVSLDSQMNLQNNHIQQQKNEKQHTPTFIFTNPWDNNEKIPQQ<br>N<br>LQYSQNYETYGMSNEDFEAFKRRKSSLVIPPARAPAPNPFHYDKYPLYS<br>N<br>INNNNINNSHRPSLGSSSSSLAPMNLQMSNPSLYQQQAQQAQDLFEKQQ<br>A<br>QQQLLSSRFLPSFYNAHNQDAQRRQSLAAPYYSQTYQSNPSVAGSKANS<br>I<br>TPSNAIPGSNSTNNYSTNAPNMTSLPAILPYRRLSAYPSSTAHLLHPSN<br>L<br>RMAATDADLLQQHLYPRPQSQYTVPSLQKCSSKSDLQPIVNPTPKFRRA<br>S<br>LHSKTISPLVGLTKNLITTYQLCSSDFTYKTSKNPKRVLTKPNEAKLNN | 119.9 |
| *C. glabrata* | CAGL0I05896<br>g | 1-336 | MVQIYYNENQALEPMQHNQANGIWKASYMDEANADSKRFAFTNPWGNEK<br>I<br>DEDEAMSYNYSSANSTADVDMNSDMYKRRKSSLVVPPSRAAAPNPFQYE<br>T<br>MQNSTGTAQRAMYPESSAYQQQRQIMVPHQQQGQLASRFNSRFVPSLYN<br>P<br>QEFQRRQSLATAQFSTPSSLHGSASNQAFNTGSGNNGVNLNYASANVTT<br>N<br>NSSTYLSAMTPNVPMEMGNIPLASPHRRLSAFPTSGTTNSSLQPPFKQL<br>R<br>RDEGSPRQIVIPQMHRCHSKNDLQPCINPNPKFRRASLHSNTISPLIGL<br>T<br>KSLLTTYALCSPDFTYQTSKNPKRVLTKPSEGKYNN | 116.7 |
| *S. uvarum* | 6.221 | 1-281 | MSQTSQRPPQQQQQHSQQQQNPQFSFVNPWGEEKITNSQQNLVYPPQYD<br>D<br>PNTNESLDAYRRRKSSLVVPPTRAPAPNPFEYDSYPAYTNSNTSLPANG<br>Q<br>FSSAYQQQQQQQVYQQSTIHPSQFGSRFVPSLYDRQEFQRRQSLATTNY<br>S<br>SNFPSINSNANQGTSSIPAMSPYRRLSAYPPSTSPPLQPPFKQLRRDEI | 105.8 |

| | | | | |
|---|---|---|---|---|
| | | | Q<br>GQKLSIPQMQPCNSKNDLQPVINATPKFRRASLNSKTISPLVSVTKSLI<br>T<br>TYSLCSPDFTYQTSKNPKRVLTKPSEGKCNN | |
| *S. kudriavzevii* | 10.73 | 1-332 | MNTSNNNDSTSSNGKNTSLSPTLATHSDASMGSGGASQDTSHLGSSIWN<br>P<br>SYMNQSSQRPLQKQQQLQQQQNPQFCFVNPWNEEKVTNSQQNLVYPPQY<br>D<br>DLNTSESLDAYRRRKSSLVVPPTRAPAPNPFQYDSYPAYTSSNTSLPAN<br>N<br>GGQYPFAYQQQQHAYQQGAIPPSQYGTRFVPSLYDRQEFQRRQSLAATN<br>Y<br>SSNFSSVNSNANQGTSSIPTISPYRRLSAYPPSTSPPLQPPFKQLRRDE<br>V<br>QAQKLSIPQMQPCNSKSDLQPVLNATPKFRRASLNSKTISPLVSVTKSL<br>I<br>TTYSLCSPDFTYQTSKNPKRVLTKPSEGKCNN | 117.6 |
| *S. mikatae* | 10.91 | 1-339 | MNTSNNNDSTSTNSNKNASLSPTVATNSDASVGSGRASQDNSHLGSSIW<br>N<br>PSYVNQSSQRHPQQQQQQQQNSQFCFVNPWNEEKVTNSQQNLVYPLQY<br>D<br>DLNSNESLDAYRRRKSSLVVPPARAPAPNPFQYDSYPAYTSSNTNLPGN<br>S<br>SGQYPSAYQQQQQQQRQQQHAYQQGTIPPSQFGSRFVPSLYDRQEFQRR<br>Q<br>SLAATNYSSNFSTFNSNANQGTSSIPVISPYRRLSAYPPSTSPPLQPPF<br>K<br>QLRRDEVQAQKLSIPQMQPCSSKNDLQPVSNATPKFRRASLNSKTISPL<br>I<br>SVTKSLITTYSLCSPDFTYQTSKNPKRVLTKPSEGKCNK | 115.2 |
| *Z. rouxii* | ZYRO0D0297<br>0g | 1-335 | MWNPLYLSSEDQEQLKKQQQQQGLQQRLHQMERRQSQSQRAQHQQHSQQ<br>H<br>QQPQQQVQFTFTNPWGNNEDLDPYYSSSAGASSTGLDTSETNPLLAAEY<br>G<br>GDGFDTYRRKSSLVIPPARAPNPNPFEYDDRNAYAGVYDPQRQNQIRNM<br>L<br>LAQQQQQLAGARFVPSSFYGAGGDLHRRQSVAAVHYPPSGPTNLASLGA<br>P<br>TNFTPAAGAATNMTPYRRLSCYPPSTSPTASLQPPYKQLRRDAGAQGAM<br>V<br>GAPSPQQVKIPQMQKCNSKSELKPTLNATPKYRRASLNSKTISPVIALT<br>K<br>GLITTYSLCSPDFSYQTSKNPKRVLTKPNEGKYNG | 117.3 |
| *T. delbrueckii* | TDEL0H0089<br>0 | 1-303 | MQQGAEATGNEEANNGNQSIWNPLFMSQEENVQQLPPPQGRQQVQFNFV<br>N<br>PWGVANGENAQVNQDNVPDETAYLTPSDKENVDSFARRKSSLVIPPARA<br>P<br>GPNPFEYDDAKAYPNMYSQRQRPMFFPQQQQQLPQQTYNSGRFVPSGFY<br>N<br>PQDSQRRQSVAVVHYPTTAPNTSNASTMIPATGATHMNSPNRRLSAYPP<br>S<br>TSPTTSLQPPYKQLRRDQGTAPPQQIIIPQMQRCTSKTDLSPMVNATPK<br>F<br>RRASLNSTTMSPLIALTKNLITTYSLCSPDFSYQTSKNPKRVLTKPSEA<br>K<br>CNS | 117.0 |
| *K. lactis* | KLLA0A05819<br>g | 1-307 | MGDKNSLWNPAFMASQQGASSSNQNQNFSQTQANTQNQGPEIAPDGPAG<br>S<br>SSNAGGNIYAGQQQGGNAAGGFQMQSPRQSIFKNPWQEIPQIPESQRST<br>M<br>RKQSNFPLPTTYEEDNAVDESSQQFRRRNSSLVIPPARAAGPDPFLYEK<br>Q<br>PFPNYNDAQRRQSVAVTGGSQLSPQQVTKPSQLRKPGINTAGYMAGSIN | 121.0 |

| | | | | |
|---|---|---|---|---|
| | | | S<br>SVYPATAGAYMSTRGGSSSLPSSRPPSPIIIVPKFNKIFTRQDLNPVIH<br>S<br>TPKYRRASLSSKTVSPLMALTKSLTTTYSLCNDDFAYQTSKNPKRVLTK<br>P<br>NEGKYNN | |
| *E. gossypii* | ACR249C | 1-248 | MKEERKSIWNPAFDMSAGAAKTGKVLGRQSFRNPWGEPGSPAGRRQSGF<br>Q<br>QLATTFEEGQGEEERARRRSSLIVPPTRAAGPEPLLRETYGDSWGYGSE<br>F<br>QRRQSVAVAGTYHSPGYFETGAQQPSTSPSLMVAHAVPYRKLSAYPPLA<br>G<br>ALVPAASLNSTLRGSAGGVAAAVPQMRKVGARQELAPVLHAMPKFRRAS<br>L<br>NSKTVSPLIALTKSLIITYSLCSDEFSYQTSKNPKRLLTKPSEGKLNN | 110.6 |
| *L. kluyveri* | SAKL0C0624<br>8g | 1-332 | MNNPVGKAQEDGTTQFQSQLPPQPLQEPQQQPQQQPQQQPQQQPQQQPQ<br>Q<br>QPQQQPQQQSIWNPAFMSNASGSTTPQGQQTFPNPWTDSTNSIHASPAQ<br>R<br>RLSGFNQQLPTTYEAVQADSQSNFRRRNSSLVIPPTRAAGPDPFLYDAQ<br>Q<br>QQQQQIFPFYHQLQQYSINQDAQRRQSVAVAGSYNRQSMGYLTEGTNYM<br>I<br>QQPQQLLQQQQQQQPQQPQQGHSPYRRLSAYPVTAGTVLPPPFKQIRR<br>D<br>SSTPVAIPRLHRVSARQDLRPVINATPKHRRASLNSKTVSPLVALTTSL<br>T<br>TTYTLCSPDFSYQTSKNPKRVLTKPSEPKYNN | 110.0 |
| *L. thermotolerans* | KLTH0F05522<br>g | 1-250 | MPESDKSIWNPAFLNNAQKQQGAPSYGFKNPWHDVSGATGKRNSMQFNQ<br>H<br>LPTTYEEAPQHGSASENGFRRRNSSLIVPPPRAAGPAPDAYMYGMQFVP<br>Q<br>QGYPGNSQELHRRQSVAVAPQHSFAQPALPADMSMHAPMSPFHRKLSSY<br>P<br>ATAGSVLPPPVKQMRRQDEMIPVVLPQMHKVNARQDMRPTINATPKYRR<br>A<br>SLDSRTVSPLVALTKSLTTTYTLCSPEFSYQTSKNPKRVLTKPSEGKNN<br>N | 109.3 |
| *L. waltii* | 33.13984 | 1-253 | MPDSDKSIWNPAFLSNSQKQPASPAYGFKNPWNDATSAAAKRNSMQFSQ<br>Q<br>LPTTYEDIQHNAGMSENEFRRRNSSLVVPPPRAAGPGPDAYMYGMQFVP<br>P<br>QGYPPYAQELHRRQSVAVAPQHTQHAFTQPSSAVDHPMHAPMSPFHRKL<br>S<br>SYPPVAGSVLPPPVKQLRRQEEVMPVVLPQMHKVNTRQDLRPAINATPK<br>Y<br>RRASLNSKTVSPLVALTKSLTTTYALCSPDFSYQTSKNPKRVLTKPSEG<br>K<br>HNN | 110.1 |

**Table S3. Yak1 homologs.** Yak1 homolog IDR sequences, along with organism, reference ID, residue positions, and ALBATROSS-predicted $R_e$.

| Organism | ID | Res. # | Sequence | Pred. $R_e$ (Å) |
|---|---|---|---|---|
| *S. cerevisiae* | YER161C | 57-273 | QELLKNGALAKKSGVKRKRGTSSGSEKKKIERNDDDEGGLGIRFKRSIG<br>A<br>SHAPLKPVVRKKPEPIKKMSFEELMKQAENNEKQPPKVKSSEPVTKERP | 99.6 |

| | | | | |
|---|---|---|---|---|
| | | | H<br>FNKPGFKSSKRPQKKASPGATLRGVSSGGNSIKSSDSPKPVKLNLPTNG<br>F<br>AQPNRRLKEKLESRKQKSRYQDDYDEEDNDMDDFIEDDEDEGYHSKSKH<br>S<br>NGPGYDRDEIWAMFNRG | |
| *V. polyspora* | 1025.5 | 58-288 | QELIKSGKLNPKGTSNKSRSSSSTPGGKRKNSPSDNDNGTEFKFKRKIN<br>S<br>NNLPKFQKPVETKHAPLKKMSFDELMKQAENNAHSKPESEPVLNSKSKE<br>L<br>NVGKSVQQKYRISKQGFKSNRNERMHNSTSVRNSRPSPKPHTTSHDLKP<br>V<br>IVPIPKGGGLAKPNEKLRERLEMKKQKLRRGRYEDEEEDEDDYDDDMDD<br>F<br>IDDDEEDYSSVSRSHKANYNRDEIWAMFNKG | 98.3 |
| *T. phaffii* | TPHA0B03460 | 58-282 | KENLKNNITKNKAASRSATLTRQRSSTTSKIDNSTETTFKRKPGQNTKA<br>F<br>SNQGQLKKKPTALKKISFEDLMKQAESNNVPNSNGDQSLKNVNNKRPLL<br>T<br>KPGFKSRKVTKPNMKNVRRSEVSVTHRAENISKKDNGPVMVKLPTMGIA<br>K<br>PNAKLIQKMKHKNSKGKGFGDRYGKSSQDHYDSEEDSDLDDFIDDDEDN<br>D<br>GYGDLEAAGDPGYNRDDIWAIFNKG | 104.3 |
| *T. blattae* | TBLA0F03750 | 55-293 | KELLKNGGDVHKKSAAKKESSLKSSTTKSSSRRPKNRDSHSETTYKRKI<br>G<br>ERTKGASYQGSNNVISRKEHTKKMSFDELMKQAETNKTKPEIDMNKQNL<br>P<br>KPARRLSKPGFKPSKYARNNQIAKANSTDLNSERKSHNSNNLRDTPSLS<br>K<br>PSGEESPVVVQIPKNNFARPNEKIRKMLDSRKRSHKRQYDYDDEEEDDM<br>S<br>DFIEDDEGEEDSYHNYDRRKADRDPGYDRDEIWAMFNKG | 105.3 |
| *N. dairenensis* | NDAI0C06050 | 60-288 | QERLKNGELEKKKPKPRRSPGSSSGSKSTRKRDNDDVEGSLGTVYKKKV<br>G<br>SSNAKLVPVRSLTKKLEPIKKLSFEELMKQAESNSKGGSPDNTAVKVTA<br>S<br>IKKTRPPPRISKPGFKSFKERKTTASKSTSTVNNKSKPEFSRNHPNGRH<br>L<br>KEQSAVKLKIPKNIVAQPNRMIKQKLESKRRTLESKYSNRRGRYEDQYD<br>D<br>DMDDFIEDDEEEEEENYRSKSRRDERPWL | 106.8 |
| *N. castellii* | NCAS0A03730 | 63-292 | KEQIKNGEFAKKHKKTNPSTTTKRKSKKDDDNLAADGYSRFKKKLGSTH<br>T<br>RPTPVRTLTRKMEPIKKISFDELMKQAENNASSKESSEGISKKESPSAS<br>R<br>PHLHKPGFRSARDRNRVSKPVKHQTTLPRKKMSLSPIRNRPGSRDATPI<br>K<br>ISLPVAQPNQRLKQRLESKRQRPSGRDRYGRPEYDYDDEDDMDDFIEDD<br>E<br>EDSEVHRRMKLHRDDPGYDRDEIWAMFNKG | 105.2 |
| *K. naganishii* | KNAG0G01760 | 61-247 | KERLKSGPTGTAVKRKRRAPGAPDEPRKRRNGADSEGGLLGTVYKRRPG<br>S<br>RQQQAATHAGAATKRDAVKKMTFEELMMQAETNAVEKPATTAPPQRTAA<br>P<br>VRNPGFKPRRRRTGPASTTKTPAEPSKPTQRRPATPRFPAQPNDLLRRR<br>L<br>KQREQREQRQQQQQHRGATAATTPRRTLSWTTSSRT | 107.1 |
| *K. africana* | KAFR0B02460 | 58-265 | KEQAINNVISKKPRARKRPSSGPKNKVAKEDGGDIGTVYKKKIGSNTTV<br>S<br>RPQVKKPAPLKKMSFEELMKQAENNATISPTVKSEAKHETSSANRIMKP | 108.3 |

| | | | | |
|---|---|---|---|---|
| | | | N<br>FKHSSSKLNPRHKIDARAKIEPGKEKPVRLSLPKNKFAQPNDRIRKELE<br>S<br>RKKHKQGYRRNELDEEDSDLSDFIEHSDDDDRYKRRTLTSYDDPGYDRD<br>E<br>IWAMFNRG | |
| *G. glabrata* | CAGL0L11704g | 55-270 | QELLKNPELAKKKQKQVRKTPSSSKASTGKKDKNGDDNMLVSRFKRKVG<br>S<br>DKPAVPIQVKKKPQPIKKLSFEELMKQAENNQTIPVSKDTQSNGAGEKI<br>K<br>GSAKLNKPGFKTSRPKSLSPTTHINKTDHGKDKSTAKEKSEPVVKIGIP<br>K<br>FAQPNERLKKKLEMRQRVNKSRRYEDEEDDMDDFIEDDEEEYSSYRTTS<br>K<br>DPGYDRDEIWAMFNKG | 107.9 |
| *S. uvarum* | 5.295 | 57-275 | QELLKNAALAKKNGVKRKRGTSSGSEKKRKERNDEDEGGLGIRFKRSIG<br>A<br>SHAPLTPAVRKKPEPVKKMSFEELMKQAESNEKQPIKTKSPEPVSMERP<br>R<br>LNKPGFKSSKKPLKKASPGLASRETPSRGDNMKLAEQHKPVKLNLPTNG<br>F<br>AQPNRRLKEKLDSRKQRSRYQEDYYEEDNDMDDFIEDDEDEEEAHRSRS<br>K<br>HTDGPGYDRDEIWAMFNRG | 99.9 |
| *S. kudriavzevii* | 5.300 | 57-273 | QELLKNGALAKKSGVKRKRSTSSGSEKKRSERNDEDEGGLGIRFKRSIG<br>A<br>SHAPLKPVVRKKPAPIKKISFEELMKQAENNEKQPAKVKSPEPIAKVRP<br>H<br>LSKPGFKSSKRLQKKPSPGTTLHGTPSRDNGVKSPESPRPVRLNLPTNG<br>F<br>AQPNKALKEKLESRKQKSRYQDGYYEEDNDMDDFIEDDEDESYRRRSKH<br>G<br>SEPGYDRDEIWAMFNRG | 104.3 |
| *S. mikatae* | 5.333 | 57-273 | QELLKNGALAKKSGVKRKRNNLSGSEKIKAERNDDDEGGLGIRFKRSIG<br>A<br>SHAPLKPVMRKKPEPIKKMSFEELMKQAENNEKQPSKVMSPEPVVKERP<br>H<br>FNKPGFKSSRRPQKKLSPSTPLRGTPSKDKGMKLSESPKPVRLNLPTNG<br>L<br>AQPNRRLKEKLDSKRQKSRYQDNYYEEDNDMDDFIEDDEDEGDHRRSKH<br>N<br>NGPGYDRDEIWAMFNRG | 101.5 |
| *Z. rouxii* | ZYRO0B11550g | 58-284 | QELLKKGELSKKATASRRTSASGKGTKKDSDDDNKTVTRFKKKSSSGSS<br>T<br>GSNRPTHISVAQKKKPEPVKKMSFDELMKQAENNAHSNGSSSSKSGSTT<br>P<br>VRAPSETQPRPRPHINKPGFKNPDRRRRAGSQEPVKPVVKNQPTPTVKE<br>P<br>RPAKLSLPKKDFAKPNELIRRRLEAKKNASRVQETQQDDYESDMDDFIE<br>D<br>DEEEDRVAMEKDPGYDRDEIWALFNRG | 108.2 |
| *T. delbrueckii* | TDEL0A01380 | 58-277 | QELLKSGELSKKVRGQSKPAPTRRRKDDDEGGSMGTKFKRKVRPHSAGS<br>S<br>LPTTNAKKAEPLKKLSFDELMKQAENNAQEKPNAMKEPSPVNSGPHRIP<br>K<br>SQGTTQQYVKKIGFKKGADRRNRNSTSPVPEIPTTKPYRDEPAPIKLKT<br>M<br>VNGFAKPNEKLRRQLEQRKRTIKPRTEYEDDGSDLDDFIEDDTMESENR<br>R<br>QSQRDTGYDRDEIWAMFNRG | 106.0 |
| *K. lactis* | KLLA0F14487g | 58-244 | LKQGISSTSTKPAAKARHKASSPKDEAPIYKKKPGVNTTSGKYPVVVPK | 102.7 |

| | | | | |
|---|---|---|---|---|
| | | | R<br>EPIKKLSFDELMKRAEQKSREGPKEDKKPLPKKLGFDKAAKPKAATKDA<br>K<br>PVKEPKSKVMVKSFNRFAQPNEKLAKKLKLKEKKQIMARHGDQHYDSEN<br>D<br>EDLSDFIEDDDLDECEQGSKSQPYDRDEIWSIFNKG | |
| *E. gossypii* | AGR161C | 62-277 | QEQLRKGTLKKASSQRRSKANGGEVASGGVRQEGSTRWKLPRPKSTVVA<br>A<br>AAPAPPLKKLSFEELMKQAEEKAKSPAASGKRTAPAGPSAPAVSKPGFK<br>P<br>RSNSGAAVVGGKVAGADKGARNGGADRTAHAPNSARGMKAKQAIAIDLP<br>S<br>GGGLAKPNEKLRRILEKQERRKRSAGEYEEDDSDLDDFIADDDGEEEGG<br>S<br>YGYDKEEIWSIFNKG | 105.7 |
| *E. cymbalariae* | 7222 | 62-268 | QDQIKRGSLGTPRSGSRKRLKTPGASRSNKGELSEEDDKVSFTTKWKLP<br>N<br>PSKVSKPSVPKGPLKKLSFDDLMKQAEEKAKSSKPEPDVATQSSRTTTT<br>K<br>LTKRGFKDKTRHTPKALVKGFTGGSPVKKKPIEKPVKIRVPSSNGIAKP<br>N<br>EKLRKMLEKRNIKRQTTEFEGNGSDLEDFIDDEEDEVDSQDGYGYNKDE<br>I<br>WSIFNKG | 106.2 |
| *L. kluyveri* | SAKL0H18084g | 66-263 | KEQLKKGTLQTKKPSVSRKRLDDTVTEARFKRKVRSSTVTRQTLPVNRQ<br>P<br>IKKLSFDELMKQAEQKSKNPPLSPSPLSKNEKKKDVKPVGLSSKIRKNG<br>F<br>KLPHQKRPVTNAKKNVPVSKKEVPVKIALPKNNIAQPSEKLRKRLEMKQ<br>Q<br>HRKRYYGDEDEDEDDMDDFIDDDDDEEAEYVKDHGYDRDEIWAMFNKG | 97.8 |
| *L. thermotolerans* | KLTH0G14586g | 66-258 | KEQLKNAPKNKPGAPSSRKKKDENSATATETKFRRKVGQSTKPQRPVAP<br>V<br>RREPLKKLSFDELMKEAEKKASDPSTDSKAPSAQKATSSIAKPIKLNKP<br>G<br>FNKGARRTPAAAPVSKPRERKEPTVKLKQLSIPKSSIAQPGEKLRKKLD<br>S<br>IKKKRQGEAYGYEDEEDLDDFIEDDEEEEGFNRDEIWAIFNKG | 101.7 |
| *L. waltii* | 56.23413 | 67-257 | KEQLKNAPKNKPAAPSRKRKDENSANTETKFRRKVGESLQSRKPVAPVK<br>R<br>TPLKKLSFDELMKEAEEKSKNPSTDPIDSTSRSKALQNNPPVRLQRPGF<br>K<br>SAARRDRKPLSTPKITKQKSPVESLQRLPAPRPSIAQPGAKLRKRLENL<br>K<br>KHRQTDRYRSSEEEDMDDFIEDDEEEQGFNRDEIWAMFNKG | 100.9 |

**Table S4. Spt2 homologs.** Spt2 homolog IDR sequences, along with organism, reference ID, residue positions, and ALBATROSS-predicted $R_e$.
